# *In vivo* suppression of polyglutamine aggregation via co-condensation of the molecular chaperone DNAJB6

**DOI:** 10.1101/2022.08.23.504914

**Authors:** Eduardo Preusser de Mattos, Maiara Kolbe Musskopf, Steven Bergink, Harm H. Kampinga

## Abstract

Amyloidogenic protein aggregation is a hallmark of several human neurodegenerative conditions, including Alzheimer’s, Parkinson’s, and Huntington’s disease (HD). Mutations and/or environmental stresses trigger conformational transition of specific proteins to amyloids, conferring them with gain of toxic function, which eventually leads to cell death in distinct brain areas. Cumulative data indicate that modulation of specific molecular chaperones can alleviate many of the pathological features of protein aggregation diseases. We previously showed that the Hsp70 co-chaperone DNAJB6 is among the strongest suppressors of amyloid aggregation, and that moderate DNAJB6 overexpression significantly extents lifespan of a mouse model of aggressive HD pathology. DNAJB6 alone delays amyloidogenic aggregation *in vitro* by several orders of magnitude at substoichiometric ratios, but its activity in cells is less efficient, albeit still markedly superior to most known anti-amyloidogenic agents. This suggests that downstream PQC factors are necessary for full DNAJB6-mediated suppression of aggregation *in vivo*, which might have to be co-stimulated in therapeutic strategies targeting DNAJB6 action. We explored here the PQC pathways required for optimal DNAJB6 inhibition of polyglutamine (polyQ) aggregation, focusing on the two main cellular proteolytic machineries: proteasomes and macroautophagy. Unexpectedly, DNAJB6 activity was largely insensitive to chemical blockage of either degradative pathway. Instead, live cell imaging unveiled a co-condensation mechanism of DNAJB6 with mobile polyQ assemblies. DNAJB6 was not required for polyQ condensation, but its expression increased the percentage of cells with mobile condensates by a factor of 3, suggesting that DNAJB6 prevents polyQ condensates to convert from the soluble to the solid state. This in turn, may keep the polyQ peptides competent for (regulated) degradation and accessible to factors allowing its extraction from the condensed state.

## Introduction

Several human neurodegenerative conditions, including Huntington’s (HD), Alzheimer’s (AD), and Parkinson’s (PD) diseases course with amyloidogenic protein aggregation and neuronal death ^1^. In the case of HD and many forms of spinocerebellar ataxia, mutations in otherwise unrelated genes lead to the production of proteins with abnormally long, highly aggregation-prone polyglutamine (polyQ) tracts ^2^. Irrespective of the native folding state, proteins such as polyQ-expanded Huntingtin in HD ^3,4^, amyloid-β in AD ^5,6^ and α-synuclein in PD ^7,8^ can adopt a comparable amyloid core, which seeds aggregation and initiates a cascade of toxic events. Upon polyQ aggregation, this includes selective cellular toxicity and neuronal degeneration of specific brain areas via primarily gain of toxic function mechanisms, that include blockage of processes like axonal transport ^9,10^, nuclear-cytoplasmatic shuttling, and the sequestration of transcription factors and components of the protein quality control (PQC) network, leading to their loss of function ^11–13^.

Much research has been dedicated to characterizing and developing treatments to protein aggregation diseases, including inhibitors of amyloidogenic aggregation ^2,14^. However, there are still no relevant therapies that prevent or significantly delay disease onset and/or progression of neurodegenerative pathologies related to toxic protein aggregation. A putative therapeutic strategy in these cases is the manipulation of steady-state levels and activity of distinct molecular chaperones, aiming at suppressing primary and/or secondary nucleation events linked to amyloidogenic aggregation ^15^. As central PQC components, chaperones play a key role in recognizing unfolded, partially folded, or misfolded polypeptides, subsequently assisting in substrate (re)folding, disaggregation, and/or targeted degradation ^16,17^. Indeed, cumulative evidence indicates that several chaperones can inhibit amyloidogenic aggregation of different disease-related proteins and at least partially rescue neurodegenerative phenotypes in animal models ^18–23^, albeit with varying levels of potency and specificity ^24^.

Screening a large library of chaperones from the Hsp70/HSPA, Hsp40/DNAJ, and Hsp110/HSPH families, we have previously shown that the ubiquitously expressed Hsp70 co-chaperone DNAJB6 is a strong suppressor of polyQ aggregation ^25^. DNAJB6 not only inhibits aggregation of distinct expanded polyQ proteins ^25–27^, but also of amyloid-β ^28–30^. Independent studies have also recently suggested that insolubilization of both α-synuclein ^31,32^ and certain yeast prions ^33^ can be prevented by DNAJB6. In fact, data indicate that specific serine/threonine (S/T) residues at the C-terminus of DNAJB6 may form hydrogen bonds with the β-hairpin structures commonly formed in amyloids, thus inhibiting primary nucleation of fibrils ^27,28^. Importantly, the S/T-rich region of DNAJB6 is crucial for its suppression of aggregation activity, as progressive deletion or substitution of these residues leads to successive loss of DNAJB6 function ^27,34^.

Recombinant DNAJB6 *in vitro* strongly inhibits amyloidogenic aggregation at substoichiometric ratios without depending on any additional factors ^27–29,34^. However, in the complex cellular environment DNAJB6 requires recycling via Hsp70 for maximal suppression of aggregation ^27^. Earlier studies also pointed to a requirement of proteasomal (but not macroautophagic) degradation for DNAJB6-mediated anti-aggregation activity ^25,35^. Nevertheless, little is known on how DNAJB6 achieves powerful suppression of amyloid aggregation *in vivo*, and the downstream cellular effectors of this process remain poorly understood. Elucidating these details is essential to better understand DNAJB6 chaperone actions, which could have clinical relevance to a wide range of human diseases with an amyloid component. Here, we used a combination of biochemical and live-cell imaging techniques to characterize the specific requirements of DNAJB6 to suppress amyloidogenic aggregation in human cells.

## Results

To better understand the requirements and mechanisms underlying the mode of action by which ectopically overexpressed DNAJB6 suppresses amyloidogenic aggregation, we followed the insolubilization profile of a previously characterized ^36^ GFP-tagged exon 1 fragment of Huntingtin containing 71 glutamines (hereafter referred to as Q71-GFP) over time in both wildtype and *DNAJB6* knockout cells (*DNAJB6*^*+/+*^ and *DNAJB6*^*-/-*^, respectively) using high-content live cell imaging. In agreement with our previous stationary data ^36^, *DNAJB6*^*-/-*^ cells showed a faster kinetics of Q71-GFP puncta formation, indicative of polyQ aggregation (**Figures 1A-C**). Importantly, the number of cells that ultimately formed puncta reached a quasi-plateau after ∼72 hours that was also higher for *DNAJB6*^*-/-*^ cells, compared to wildtype counterparts. The hypersensitivity to polyQ aggregation of *DNAJB6*^*-/-*^ cells was fully rescued by co-expression with DNAJB6b and, in fact, similar in kinetics and plateau values to those found in *DNAJB6*^*+/+*^ cells similarly co-transfected.

**Figure 1.**
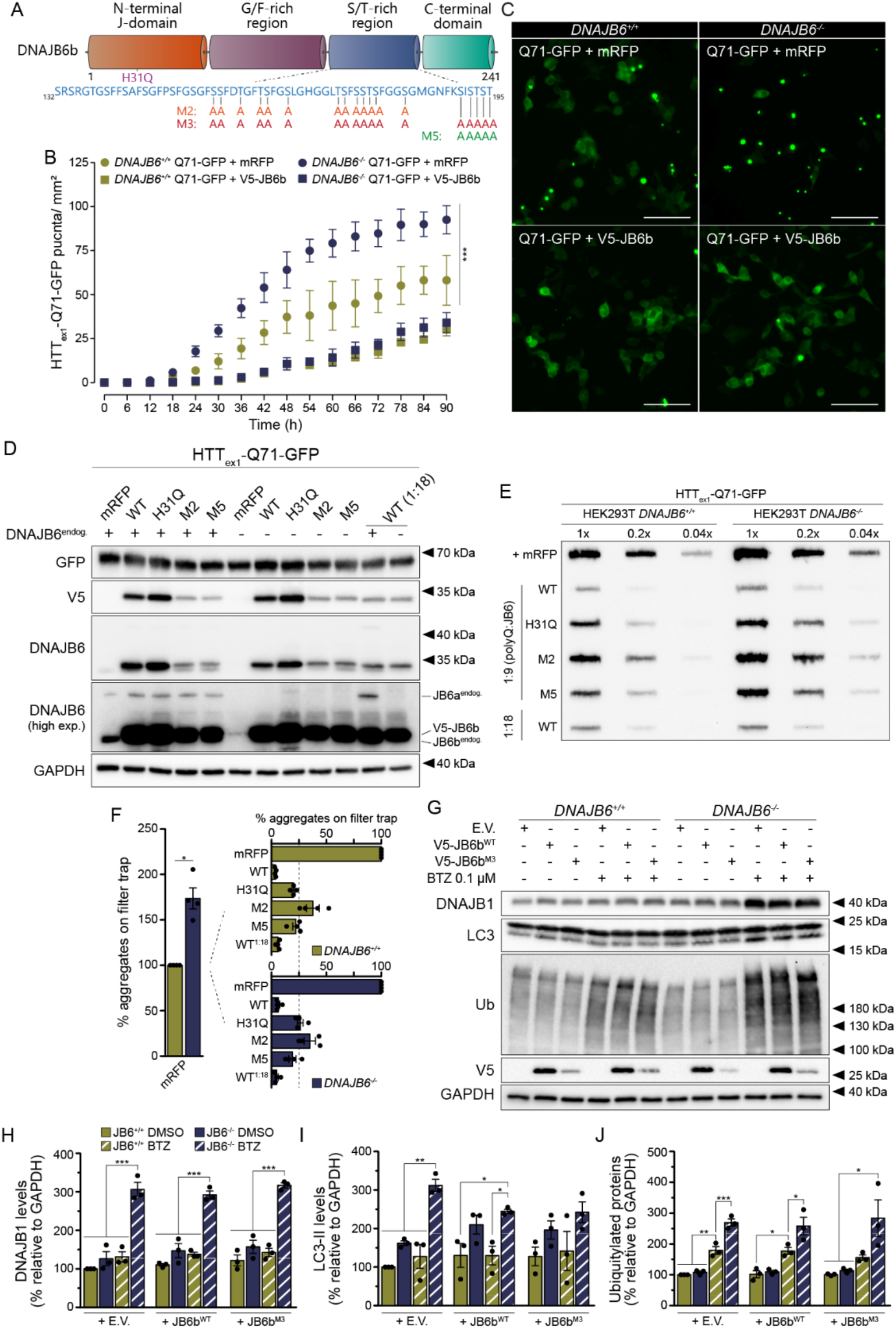
Contribution of endogenous DNAJB6 expression for full suppression of polyglutamine aggregation and maintenance of general protein homeostasis. **A:** Schematic representation and domain organization of DNAJB6b. The amino acid composition of the serine/threonine (S/T)-rich region is highlighted in blue, and alanine substitutions in mutants M2, M3, and M5 are depicted in orange, red, and green, respectively. Numbers represent amino acid positions. G/F: glycine/phenylalanine-rich region. **B:** Quantification of live cell imaging time course of Q71-GFP puncta formation in HEK293T wildtype (DNAJB6^+/+^) or DNAJB6 knockout (DNAJB6^-/-^) cells co-transfected with Q71-GFP and either mRFP or V5-DNAJB6b. *** p<0.001 (F_6,180_=16.20, sigmoidal dose-response curves with variable slope fit). **C:** Representative images from the quantification shown in B at t=48h. Scale bars: 20 μm. **D:** Representative Western blot images of cells transfected with Q71-GFP and either mRFP or V5-DNAJB6b^WT^ or selected mutants depicted in (A). DNAJB6_endog._: endogenous DNAJB6 expression (+: DNAJB6^+/+^, -: DNAJB6^-/-^). High exp.: high exposure. **E:** Representative filter trap image of the amount of Q71-GFP aggregates in cells treated as in (D). **F:** Quantification of the percentage of aggregates on filter trap, as shown in (E), for four independent experimental replicates. Left: percentage of aggregates in DNAJB6^-/-^ cells co-transfected with Q71-GFP and mRFP relative to DNAJB6^+/+^ cells (* p=0.008, one sample t-test). Right: percentage of aggregates in cells co-transfected with Q71-GFP and distinct V5-DNAJB6b variants relative to the +mRFP condition in cells from the same genotype. **G:** Representative Western blot images of levels of selected protein quality control components in cells from both genotypes upon inhibition of proteasomal activity with bortezomib (BTZ) for the last 12 hours before harvesting. E.V.: empty vector. **H-J:** Quantification of Western blot bands corresponding to DNAJB1 (**H**), LC3-II (**I**), and total ubiquitylated proteins (**J**) for three independent experimental replicates. * p<0.05, ** p<0.01, *** p<0.001 (one-way ANOVA with Sídák’s multiple comparisons test between samples transfected with the same construct). Error bars represent standard deviations (B) or standard errors of the mean (F, H-J).

Western blot analysis at 48 hours after transfection revealed that total soluble Q71-GFP levels were not markedly different between *DNAJB6*^*+/+*^ and *DNAJB6*^*-/-*^ cells, nor upon co-expression with DNAJB6b (**Figures 1D, S1A and S1B**). In line with the imaging data, however, at 48 hours after transfection *DNAJB6*^*-/-*^ cells had about 70% more Q71-GFP aggregates retained on a filter trap assay than wildtype control (**Figures 1E and 1F**). In cells from both genotypes, Q71-GFP aggregation was again rescued to a similar extent by co-expression with DNAJB6b. To further corroborate these findings, we used distinct DNAJB6b variants with either impaired Hsp70 interaction (H31Q) ^25,27^ or reduced polyQ substrate binding affinity (so-called M2 and M5; **see Figure 1A** ^27^). All these mutants have slightly impaired activity against polyQ aggregation in wildtype, *DNAJB6*^*+/+*^ cells ^25,27^. Although more aggregated material was found in *DNAJB6*^*-/-*^ cells relative to the wildtype background (**Figure S1C**), the relative protective activities of all mutants was virtually identical in *DNAJB6*^*+/+*^ and *DNAJB6*^*-/-*^ cells (**Figure 1F**).

Of note, steady-state levels of mutants M2 and M5 were significantly lower than the WT and H31Q DNAJB6b variants (**see Figure 1D**), which could at least partially explain the lower capacity of M2 and M5 to suppress polyQ aggregation, compared to WT DNAJB6b. Nevertheless, transfection of only half the usual amount of WT DNAJB6b (1: 4.5 polyQ: chaperone ratio, instead of the usual 1: 9), which yielded steady-state levels similar to M2 and M5 (**see Figure S1A**), still led to strong suppression of polyQ aggregation comparable to WT DNAJB6b at the 1:9 polyQ:chaperone ratio (**see Figure 1E**). This suggests that the loss of activity of mutants M2 and M5 is primarily due to the alanine substitutions in the S/T-rich region, rather than lower amounts of chaperone molecules available to interact with polyQ, compared to WT DNAJB6b. Overall, these data support our earlier observations ^25,27,36^ that DNAJB6b activity is tightly correlated to sensitivity to polyQ aggregation. In addition, our results show that ectopic DNAJB6 overexpression efficiently suppresses polyQ aggregation without requiring endogenous DNAJB6. Finally, DNAJB6 activity determines the threshold for cells to initiate polyQ aggregation.

*DNAJB6* is an essential developmental gene ^35,37^ and mutations interfering with DNAJB6 function cause dominantly inherited muscular dystrophy presenting with largely dysfunctional muscle cells containing protein aggregates ^38,39^. We were thus intrigued by our success in generating stable *DNAJB6* knockout cells that are viable even upon cellular insults such as polyQ aggregation. To better understand the requirements of DNAJB6 to global protein homeostasis and cellular fitness, we therefore performed a more detailed characterization of the *DNAJB6*^*-/-*^ stable cell line used here.

Lack of endogenous DNAJB6 expression led to a significantly slower growth rate, which was on average two times slower than that of *DNAJB6*^*+/+*^ cells. Intriguingly, this phenotype was not rescued by ectopic expression of DNAJB6b (**Figures S1D and S1E**), suggesting it is related to cell-adaptive features triggered by the absence of DNAJB6 during clonal selection. Western blot analysis revealed similar steady-state levels of several chaperones and other PQC components between cells from both genotypes in unstressed conditions. The only notable exception was the adaptor protein P62, whose levels were consistently lower in cells lacking endogenous DNAJB6 than in wildtype cells (**Figures S2 and S3**).

Next, we asked whether cellular features of DNAJB6^-/-^ cells would change under stress conditions, focusing first on the effect of polyQ aggregation. Expression of Q71-GFP had almost no impact on cellular growth rates, irrespective of DNAJB6 complementation or genetic background. Nevertheless, variations in growth rates became very large in *DNAJB6*^*-/-*^ cells, with some experimental replicates requiring times as high as 90 hours to reach maximal confluency (**Figures S1D and S1E)**. Whilst the latter supports the notion that *DNAJB6*^*-/-*^ cells are hypersensitive to polyQ aggregation, as well as being “at the edge” of survival, these data also demonstrate the mere insensitivity of dividing cells to the consequences of protein aggregates, which are markedly different from those observed in non-dividing cells (*e*.*g*., neurons).

Given the assumed links of DNAJB6 to proteasomal degradation ^25,35^, we next exposed cells to the proteasomal inhibitor bortezomib (BTZ) for 12 hours. Interestingly, there was marked re-wiring of PQC components in *DNAJB6*^*-/-*^ cells upon inhibition of proteasomal activity. Compared to wildtype cells, steady-state levels of lipidated LC3 and the Hsp40 co-chaperone DNAJB1 were increased in knockout cells upon BTZ treatment, and these were not rescued by complementation with WT DNAJB6b or the mutant variant M3, which does not interact with polyQ substrates ^27^ (**Figures 1G-J, S2, and S4**). Except for the higher steady-state levels of LC3-II, these changes were exclusive to the inhibition of proteasomes, since chemical blockage of lysosome-dependent pathways did not result in enhanced accumulation of DNAJB1 or ubiquitylated proteins in *DNAJB6*^*-/-*^ cells, compared to wildtype cells (**Figure S3**). Proteasomal inhibition also led to a build-up of P62 in knockout cells, suggesting that *DNAJB6*^*-/-*^ cells have either lower synthesis or faster proteasomal degradation rates of P62. In line, *DNAJB6*^*-/-*^ cells showed stark accumulation of mono- and poly-ubiquitylated proteins (**see Figures 1G and 1J**). Together, these results indicate that endogenous DNAJB6 expression plays an important role in PQC, especially in the suppression of aggregation of amyloidogenic substrates such as polyQ. Moreover, although *DNAJB6*^*-/-*^ cells are viable, genetic ablation of *DNAJB6* contributes to a global protein homeostasis imbalance and leads to adaptation mechanisms that cannot be rescued by ectopic DNAJB6b expression.

DNAJB6-mediated proteasomal degradation thus seems to be relevant at least for some endogenous proteins. Even though polyQ overexpression did not affect cell survival (**see Figure S1D**), these data raise the possibility that DNAJB6 might also influence degradation of polyQ species via proteasomes. We then sought to better dissect the PQC pathways acting downstream of DNAJB6 that may be linked to or required for its activity to suppress polyQ aggregation. Hereto, we investigated the capacity of DNAJB6b to suppress polyQ aggregation upon chemical blockage of proteasomal degradation with either MG132 or BTZ (**Figure 2A**). Because long-term inhibition of proteasomal activity is toxic, we chose to add drugs for only 6 or 12 hours (MG132 or BTZ, respectively) before harvesting cells. These relatively short exposures were sufficient to cause pronounced accumulation of ubiquitylated proteins without eliciting marked toxicity (**Figures 2C and S5A-E**). Such treatments led to modest overall increases in the amount of polyQ aggregates on filter trap in *DNAJB6*^*+/+*^ cells and slight decreased capacity of DNAJB6b to suppress aggregation, with DNAJB6b mutants displaying a proportional loss of function (**Figures 2B and 2D**).

**Figure 2.**
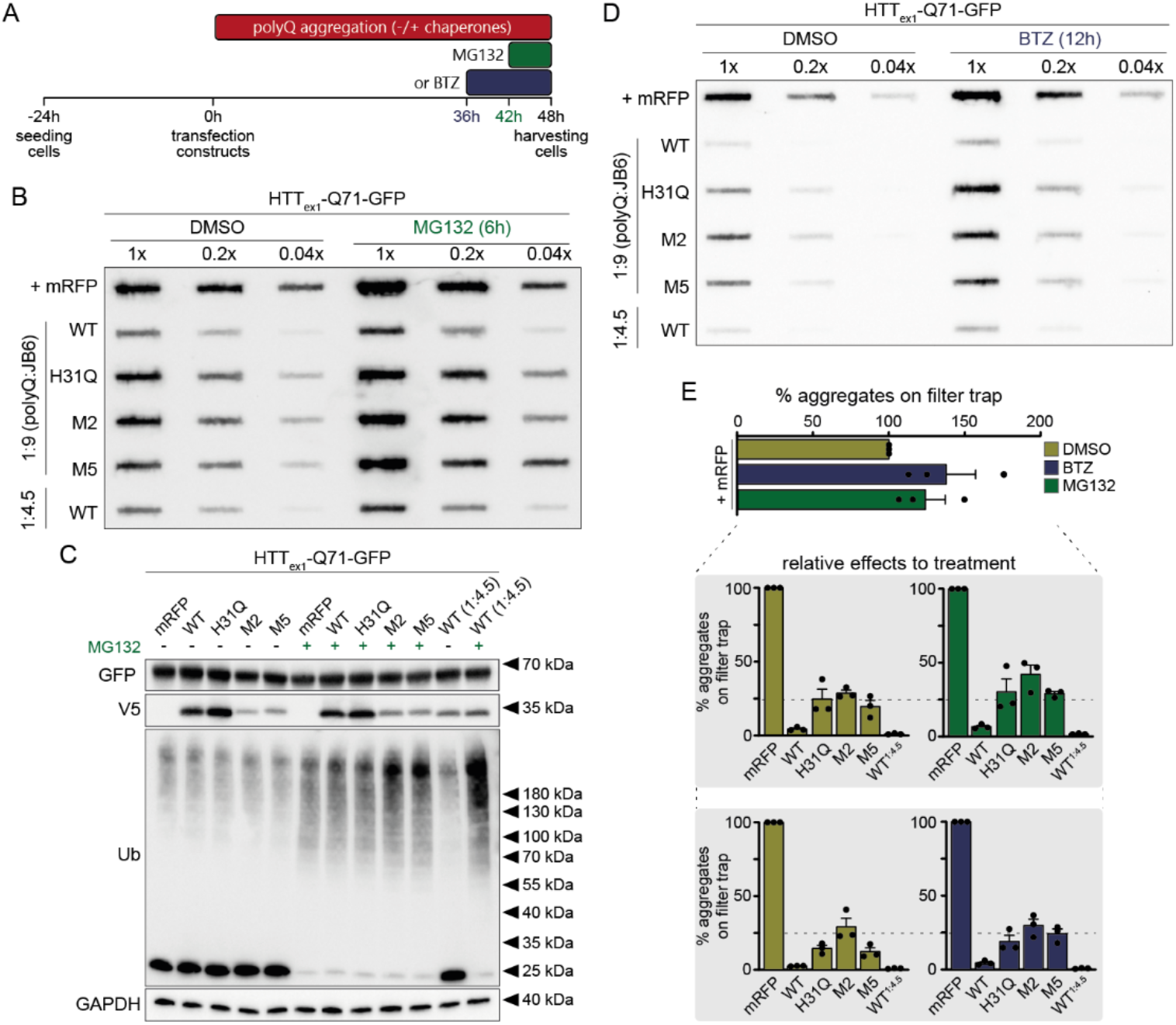
DNAJB6-mediated suppression of polyglutamine aggregation operates largely independent from proteasomal degradation pathways. **A:** Diagram illustrating the experimental regimens for the inhibition of proteasomal activity with MG132 or bortezomib (BTZ) for the last 6 or 12 hours before harvesting cells, respectively. **B:** Representative filter trap image of the amount of Q71-GFP aggregates in cells treated with DMSO or MG132 upon co-transfection with mRFP or V5-DNAJB6b variants. **C:** Representative Western blot images of the experimental conditions depicted in (B). Anti-ubiquitin (Ub) antibody was used to confirm the inhibition of proteasomal degradation. **D:** Similar to (B), but for cells treated with either DMSO or BTZ. **E:** Overview of relative effects of treatment (DMSO *vs*. MG132 or BTZ) in the amount of Q71-GFP aggregates on filter trap across three independent experimental replicates per drug treatment. Quantifications were normalized to the +mRFP control sample within each treatment group to account for the overall higher burden of polyQ aggregation upon proteasome inhibition and focus only on putative DNAJB6b loss of function when protein degradation pathways were blocked. Construct-specific differences in amount of aggregates on filter trap between the DMSO and MG132 or BTZ conditions were assessed by one-way ANOVA with Sídák’s multiple comparisons test. Data are represented as means and standard errors of the mean.

At first, these results might indicate that proteasomes are indeed required for DNAJB6b-mediated suppression of polyQ aggregation. However, it is possible that the larger fraction of aggregates on filter trap is only a reflection of the increased global PQC burden under proteasomal inhibition (*e*.*g*., diversion of the chaperone machinery to other endogenous, aggregation-prone proteins), and not a direct loss of function of DNAJB6b variants. To address that issue, we normalized raw GFP intensities on filter trap to the control condition subjected to the same treatment (+mRFP treated with either DMSO, MG132, or BTZ), thus accounting for the extra polyQ aggregation load when proteasomal activity was impaired. Remarkably, this analysis revealed that DNAJB6b-mediated suppression of aggregation for all constructs tested is unaffected by inhibition of proteasomal activity (**Figure 2E**). Hence, unlike hypothesized and hinted by the data on proteasomal hypersensitivity of *DNAJB6*^*-/-*^ cells (**see Figures 1G-J and S2-S4**), proteasomes do not play a significant role in the suppression of polyQ aggregation via DNAJB6b.

Using *ATG5*^*-/-*^ mouse embryonic fibroblasts, we have previously shown that protein degradation via (macro)autophagy was also not required for DNAJB6b-mediated suppression of polyQ aggregation ^25^. To extend these data to the hippomorphic DNAJB6b variants, we employed here a complementary approach that did not rely on knockout cell lines, but on chemical blockage of the autophagic machinery with a cocktail of three lysosome-targeting drugs (bafilomycin A, pepstatin A, and E64d) ^40^. In line with results published earlier ^25^, we confirmed that cells do not exploit the (macro)autophagy-lysosome pathway to process DNAJB6b-chaperoned polyQ assemblies (**Figures S5G-K**). Independency of proteasomal or macroautophagic degradation were also not an artifact of the timing of drug treatments relative to the polyQ aggregation process since exposure of cells to the same drugs immediately after co-transfection of Q71-GFP and DNAJB6b variants yielded essentially the same results (**Figure S6**).

To further confirm that DNAJB6b does not require active proteasomal degradation to suppress polyQ aggregation, we employed histidine pulldowns under denaturing conditions and assessed the ubiquitylation status of Q71-GFP in the absence and presence of DNAJB6b co-overexpression. Consistent with a mainly proteasome-independent pathway, DNAJB6b co-expression led to a four-fold reduction in steady-state levels of mono- and poly-ubiquitylated polyQ species, even upon blockage of proteasomal activity with MG132 (**Figures 3A** and **3B; see Figure S7** for uncropped, unedited blot). That was in stark contrast to the enhanced ubiquitylation pattern of Q71-GFP when co-transfected with E6AP/Ube3a, an ubiquitin E3 ligase previously shown to facilitate huntingtin ubiquitylation and degradation via proteasomes ^41,42^. Collectively, these data suggest that DNAJB6b-mediated suppression of polyQ aggregation in cells is related to a degradation process that is dependent on polyQ ubiquitylation, but that does not involve the proteasome or macroautophagy.

**Figure 3.**
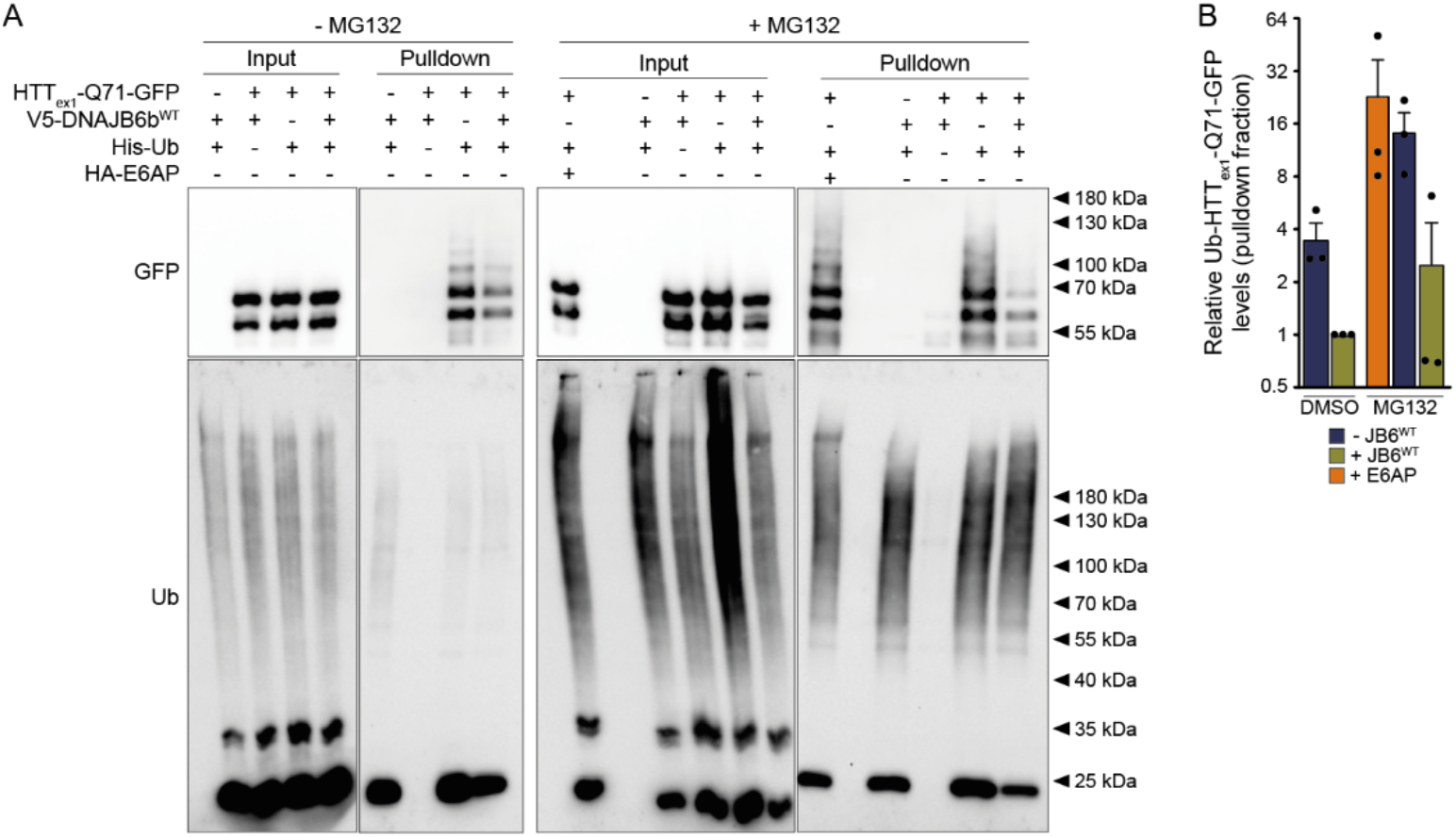
Overexpression of DNAJB6b decreases the steady-state levels of mono- and poly-ubiquitylated HTT_ex1_-Q71-GFP species. **A:** Representative Western blot images of denaturing histidine (His)-pulldowns from cells co-transfected with Q71-GFP and either mRFP or V5-DNAJB6b^WT^ in the presence of His-tagged ubiquitin (His-Ub). The ubiquitin E3 ligase E6AP was used as a positive control of ubiquitylation of Q71-GFP. **B:** Quantification of mono- and poly-ubiquitylated species in the pulldown fraction, as depicted in (A), for three independent experimental replicates (the original, uncropped version of this blot is shown in Supplementary Figure 7).

Given these elusive results, we decided to step back and ask how, and at which step, DNAJB6 affects the process of aggregation of polyQ that next might promote polyQ ubiquitylation. Our previous *in vitro* studies revealed that, even though most efficient in inhibiting primary nucleation of amyloidogenic substrates ^27–29^, DNAJB6 does not interact with monomeric amyloid species ^30^. Also, the substoichiometric effectiveness DNAJB6 actions supports interactions with non-monomeric substrates ^27,29,34^. We were therefore triggered by recent *in vitro* ^43,44^ and *in vivo* ^43,45^ studies that suggested that amyloidogenic polyQ aggregation proceeds via the formation of immiscible, liquid-like condensates which eventually transition to solid inclusions bodies (reviewed in ^46^). This process is dependent on critical threshold concentrations of polyQ species and hydrophobic interactions ^43,44^, similar to the phase separation behavior attributed to an increasing number of neurodegeneration-related proteins ^47–51^. We thus asked whether DNAJB6 could affect the balance and/or kinetics of polyQ condensation, such that it might not lead to the transition to amyloids, and hence remain accessible to ubiquitylation and further processing.

Peskett *et al*. (2018) previously used time-lapse microscopy and computational single-cell tracking to detect polyQ condensates and mature inclusions (referred to as dim and bright assemblies, respectively) in yeast and human cells ^43^. Here, we took advantage of a complementary approach based on fluorescence loss in photobleaching (FLIP) and differential exchange rates of polyQ molecules in distinct phases/ compartments ^52^. Specifically, we hypothesized that the partition of polyQ-GFP species in condensates could be more readily visualized by depleting most of the GFP signal in live cells without visible aggregates, thus revealing putative dim assemblies existing in slower exchange with the surrounding cellular environment (**Figure 4A**). To confirm the liquid-like nature of these structures, half of the condensate was then bleached with a short laser pulse, akin to fluorescence recovery after photobleaching (FRAP) experiments. Rapid loss of fluorescence throughout the whole structure, including the non-bleached half of the condensate, would thus be indicative of high internal mobility and liquid-like properties. On the other hand, if these structures are small (solid) aggregates, mobility within the assembly would be slow, and there should be significant differences in fluorescence intensity between the bleached and non-bleached segments after FRAP (compare outcomes 2a and 2b in **Figure 4A**).

**Figure 4.**
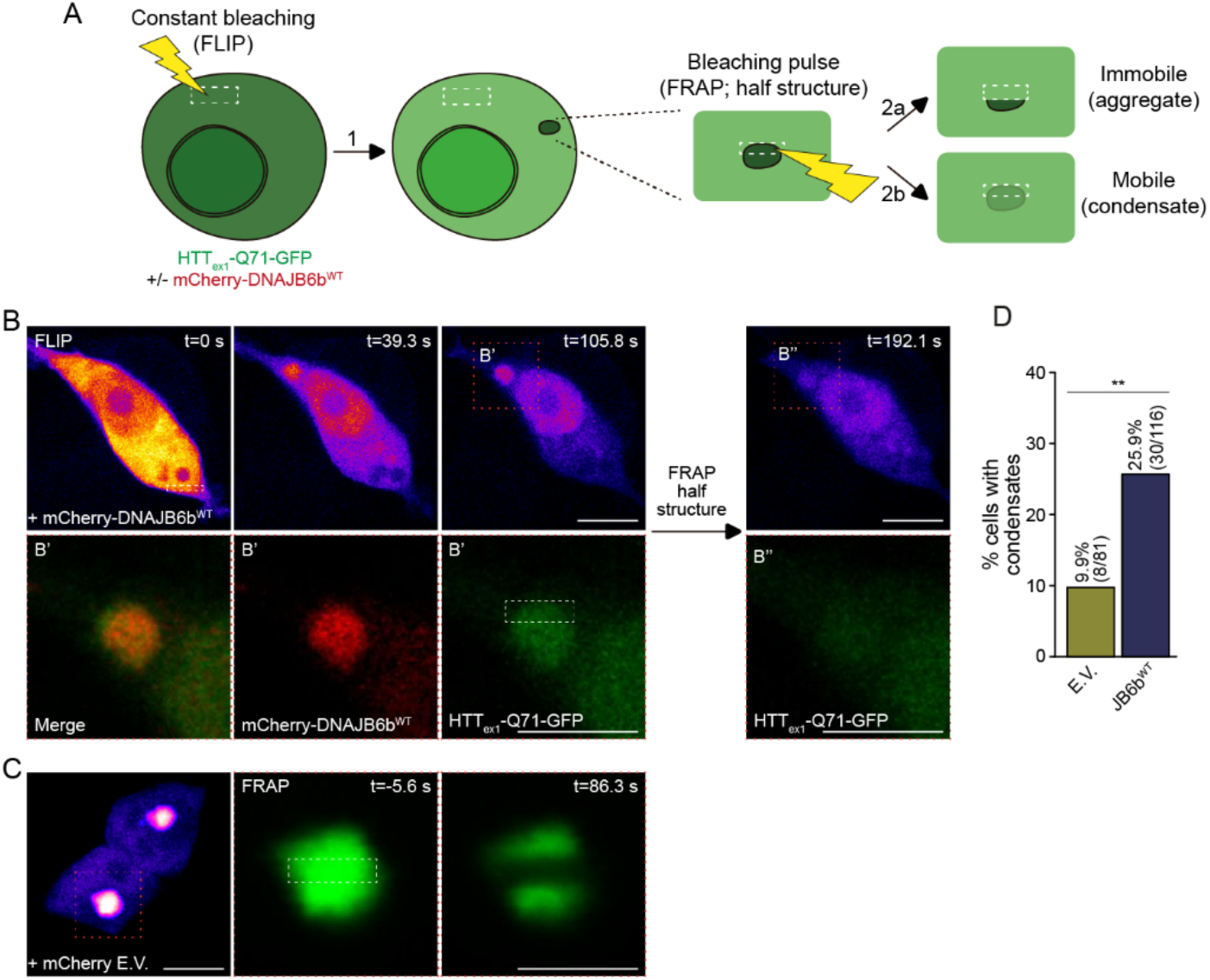
Overexpression of DNAJB6b increases the frequency of condensate-like polyglutamine assemblies in live cells. **A:** Schematic representation of the live cell imaging approach used to reveal condensate-like polyglutamine structures. Cells co-transfected with Q71-GFP and either empty mCherry vector or mCherry-DNAJB6b^WT^, and without visible aggregates, were subjected to fluorescence loss in photobleaching (FLIP), eventually leading to overall decrease in GFP intensity (1). If a condensate-like structure is present, differences in GFP intensity due to slower exchange with the surrounding environment make these structures visible. To determine whether Q71-GFP inside these structures is mobile (condensate-like), half of the structure is bleached with a short laser pulse. If the revealed structure is a solid-like aggregate, internal mobility is low, and the non-bleached half portion should retain higher fluorescence intensity (2a). If molecules inside the structure exchange freely, a rapid decrease in fluorescence across the whole structure is expected (2b). **B:** Representative images of condensate-like detection in a cell without visible aggregates and co-transfected with Q71-GFP and mCherry-DNAJB6b^WT^. Constant bleaching of a small cytoplasmic area (dashed white rectangle, upper left image) reveals a round structure of slower exchange and enriched for both Q71-GFP and DNAJB6b (highlighted in B’, dotted orange box). Bleaching the upper half portion of the assembly resulted in fast loss of GFP intensity across the whole structure, which was sustained for at least 1.5 minutes after the bleaching pulse (highlighted in B’’), indicating a high degree of mobility within the assembly [compare with polyglutamine aggregate in (C)]. **C:** Representative images of an Q71-GFP aggregate immediately before (left) and almost 1.5 minutes after bleaching its middle portion (dashed white rectangle on left) using FRAP. Scale bars in (B) and (C): 10 μm. **D:** Quantification of the percentage of cells with condensates-like assemblies, as illustrated in (B), for cells co-transfected with Q71-GFP and either empty vector (E.V.) or DNAJB6b^WT^ (N-terminal mCherry or V5 constructs; ** p=0.006, Fisher’s exact test).

To assess whether polyQ condensate formation is an intrinsic property of polyQ assemblies independent from DNAJB6, we first transfected Q71-GFP in *DNAJB6*^*-/-*^ cells and observed cytoplasmic polyQ condensates in approximately 10% (8 out of 89) of cells (**see Figure 4D**). Next, Q71-GFP was co-transfected with mCherry-DNAJB6b, and several instances of co-localization of polyQ and DNAJB6b in condensates were detected (**Figure 4B**). In all cases, there was rapid, global loss of GFP intensity within condensates after bleaching half of the structure (**Figure 4B**, inset B’’), in stark contrast with the behavior of solid polyQ aggregates, in which there was virtually no mobility detected (**Figure 4C**). Remarkably, co-expression with mCherry-DNAJB6b, or with non-fluorescent V5-DNAJB6b construct used in previous experiments (**see Figures 1-3**), led to an almost 3-fold increase in the frequency of cells with cytoplasmic polyQ condensates (**Figure 4D**; **see Figure S8** for additional examples of polyQ condensates). Together, these data suggest that DNAJB6b co-condensates with polyQ in liquid-like structures that form independently of DNAJB6, but that do not or less rapidly undergo a transition from the liquid to the solid state, thus remaining in a state that allows exchange with the cytoplasm and accessible to downstream processing.

## Discussion

Here we find that DNAJB6 partitions with condensates formed by a mutant exon 1 fragment of HTT containing 71 glutamines. PolyQ condensation occurs irrespectively of DNAJB6 expression, but the relative frequency of fluid condensates to solid inclusions is increased when DNAJB6 is co-expressed, suggesting that DNAJB6 delays the transition of condensed polyQ proteins from the liquid to a solid state. Despite the fact that DNAJB6 has been repeatedly associated with proteasomal degradation of its clients ^25,32,35,39^, we find that the ability of DNAJB6 to chaperone polyQ substrates is not dependent on proteasomal activity.

### Absence of DNAJB6 does not lead to hypersensitivity to polyQ

Maybe somewhat surprising at first sight, our data revealed that cellular growth rates are not affected by polyQ aggregates, irrespective of DNAJB6 expression or extent of polyQ aggregation. Rather than such observations being an argument against aggregate toxicity ^53,54^, we propose that this illustrates the likely very divergent impact of polyQ aggregation on replicative cells, compared to its effects on terminally differentiated, non-dividing cells like neurons. Dividing cells may tolerate such aggregates much better, for instance, because they are able to segregate the aggregates during mitosis ^55–57^, but neurons and other post-mitotic cells are not able to do so. In addition, neurons will be affected by aggregate consequences such as axonal clogging ^58^ and consequential impaired antero- and retrograde transport ^9,10,59^, and related effects such as reduced dendritic density, a result of polyQ aggregation against which a chaperone like DNAJB6 protects ^27^. This stresses that data on aggregate toxicity using replicative models may not always be informative for assessing the toxicity of polyQ proteins.

Contrary to the absence of polyQ toxicity, we found that cells stably lacking DNAJB6 expression were hypersensitive to proteasomal inhibition (**Figures 1G-J, S2, and S4**), suggesting that there might be additional, yet uncharacterized endogenous client proteins that depend on DNAJB6-mediated chaperoning for solubility, proteasomal degradation, and cell growth, even in dividing cells. Indeed, proteins like mutant PARK2 can be chaperoned for proteasomal degradation depending on DNAJB6 expression ^60^. In line, although handling of polyQ aggregates by DNAJB6 seems unrelated to proteasomal activity, expression of soluble polyQ was affected by DNAJB6 expression in a Hsp70- and proteasomal-dependent manner ^25^.

### Role and function of DNAJB6b co-condensation with polyQ

*In vitro* studies have shown that recombinant DNAJB6 efficiently inhibits amyloidogenic aggregation of polyQ but also amyloid-β at substoichiometric concentrations by several orders of magnitude ^29,34^. This implies that DNAJB6 likely does so not by interacting with monomeric peptides, but rather at a stage where they already have conglomerated ^28^. Indeed, experimental data showed *e*.*g*., that DNAJB6 has low affinity for monomeric amyloid-β ^30^, yet inhibits primary nucleation into the amyloid state ^27,29,34^. Our current data confirmed the observations of Peskett *et al*. (2018) that polyQ peptides form mobile condensates ^43^, and we now show that DNAJB6 co-associates with these assemblies. Whilst DNAJB6 is not required for polyQ condensate formation, it increases the frequency of cells with mobile polyQ condensates, while reducing the fraction of cells showing immobile inclusions. This indicates that, via co-condensation with polyQ, DNAJB6 changes the physico-chemical properties of polyQ such that it does not undergo transition to a more solid state. Considering the substoichiometric ratios at which DNAJB6 suppresses amyloidogenic aggregation, it is tempting to speculate that single molecules of DNAJB6 transiently interact with multiple polyQ peptides inside condensates, thus prohibiting amyloidogenic transition into fibrils.

### DNAJB6 and polyQ protein processing

Whilst DNAJB6 substantially reduces polyQ aggregation and toxicity in cells, the magnitude of its effect (5-to 20-fold reduction in aggregation for a tripling of its concentration ^36^) is much smaller compared to what has been observed *in vitro* (10^9^-fold inhibition of primary nucleation at a 1: 10 ratio ^27,28^). This implies that there might be cellular bottlenecks that could become rate-limiting under conditions where DNAJB6 is boosted *in vivo*, and which might have to be co-stimulated in prospective therapeutic strategies. To this end, we explored how DNAJB6-chaperoned polyQ is further processed in cells.

One possibility would be that DNAJB6 keeps polyQ condensates competent for degradation. As our previous data revealed that DNAJB6 was still capable of suppressing polyQ aggregation in ATG5^-/-^ cells (*i*.*e*., cells that cannot perform macroautophagy) ^25^, we addressed whether proteasomes would act in the downstream processing of DNAJB6-chaperoned polyQ substrates. This was also instigated by our finding that polyQ steady-state levels were dependent on expression of DNAJB6 and its ability to interact with Hsp70 ^25^. Nevertheless, DNAJB6 anti-aggregation activity was found to be largely insensitive to prolonged inhibition of the UPS, in line with previous results ^25^. However, the holdase capacity of wildtype DNAJB6 towards amyloidogenic clients is remarkably high, and one could argue that proteasomal dependency might not be revealed under these conditions. In fact, DNAJB6 dependence on Hsp70 was only revealed when we combined the DNAJB6 H31Q variant (which does not interact with Hsp70) with S/T-rich region mutants that behave as hypomorphs for holdase activity ^27^. So, to be able to more conclusively exclude proteasomal involvement, we also analyzed here these hypomorphs and still found that proteasomal inhibition did not lead to further loss of their (reduced) anti-aggregation activity (**Figures 2, S5, S6**, and references ^25,27^). So, differently from the DNAJB6-dependent proteasomal degradation of soluble mutant HTT ^25^, polyQ-DNAJB6 condensates do not seem to be a substrate of the ubiquitin-proteasome system.

But what then is precisely the fate of polyQ species in the condensed phase when DNAJB6 is present? Our current data with chemical inhibition of macroautophagy combined with hypomorphic DNAJB6 variants confirmed that this proteolytic pathway is also not involved. It is possible that inside condensates, where polyQ concentration is high, DNAJB6 prevents amyloidogenic transition, while also allowing the release of polyQ peptides from condensates to the cytosol, where polyQ concentration is too low to favor aggregation. Such dissolution of polyQ species from condensates could either be passive or due to active processes (*e*.*g*., Hsp70-dependent). In fact, co-condensation of DNAJB6 may be crucial for the recruitment of Hsp70 to polyQ condensates for further processing, as suggested by recent work from Klaips *et al*. (2020) ^61^. Recent studies investigating phase separation of the RNA binding protein TDP-43 also corroborate this hypothesis, with data showing that Hsp70 activity is crucial to maintain the liquid-like properties of de-mixed TDP-43, thus keeping it in a degradation-competent state ^49^. Similarly, extraction of aggregation-prone polypeptides from phase-separated assemblies inside nucleoli after heat stress requires refolding via Hsp70 ^62^.

In the case of DNAJB6-mediated suppression of aggregation, it remains to be established (a) whether single polyQ peptides or oligomers are released from condensates and (b) what the fate of the material released in the cytosol would be and whether non-proteasomal and non-macroautophagic pathways are involved in further processing. Nevertheless, given that mechanisms ensuring PQC and protein homeostasis decline with aging and in disease states ^17,63,64^, it is promising that substantial suppression of aggregation might be achieved (provided sufficient DNAJB6 levels) even in cell populations with impaired protein degradation.

## Acknowledgements

The authors would like to acknowledge the following funding agencies and grant sponsors: Nederlandse organisatie voor wetenschappelijk (NWO)-Top (Netherlands), Campagneteam Huntington (Netherlands), European Union Join Program – Neurodegenerative Disease Research (JPND), Science Without Borders Program (Ministry of Education, Brazil), and Graduate School of Medical Sciences, University of Groningen (Netherlands). Part of the work has been performed at the University Medical Center Groningen (UMCG) Imaging and Microscopy Center (UMIC), which is sponsored by NWO grant 175-010-2009-023.

## Materials and Methods

### Cell culture, constructs, and chemicals

HEK293T wildtype (American Type Culture Collection, CRL-3216) and DNAJB6 knockout cells ^36^ were cultured in Dulbecco’s Modified Eagle Medium (Gibco, Dublin, Ireland) supplemented with 10% fetal calf serum, 100 U ml^-1^ penicillin and 100 μg ml^-1^ streptomycin (Invitrogen, Carlsbad, USA), and 1x GlutaMAX (Gibco), and kept at 37ºC with controlled humidity and 5% CO_2_. The DNA constructs used in this study are described in **Table S1**. pcDNA5 FRT-TO mCherry was generated by subcloning an mCherry plus stop codon fragment from pcDNA3.1(+) mCherry (kind gift from B. Giepmans, University Medical Center Groningen, Netherlands) using XhoI/ NotI restriction sites. pcDNA5 FRT-TO mCherry-DNAJB6b^WT^ was generated by amplification of DNAJB6b^WT^ from pcDNA5 FRT-TO V5-DNAJB6b^WT^ and cloning into pcDNA5 FRT-TO mCherry using XhoI/ BclI restriction sites, followed by removal of the mCherry stop codon by site-directed mutagenesis. All chemicals were purchased from Sigma-Aldrich (St. Louis, USA) unless otherwise stated. The following drugs (final concentration) were used for inhibition of proteasomal or lysosomal activity: bortezomib (0.1 μM, Selleck Chemicals, Houston, USA), MG132 (5 μM), bafilomycin A (0.1 μM), pepstatin A (14.6 μM), E64d (10 μM).

### Western blot analysis

Unless otherwise stated, cells grown on 6-wells plates were scrapped on ice with 200 μl lysis buffer [10 mM Tris-HCl pH 8.0, 150 mM NaCl, 2% sodium dodecyl sulphate (SDS)], collected into 1.5 ml tubes, and sonicated on ice for 5 s at 50 V. Protein concentration was estimated with the DC protein assay (Bio-Rad, Hercules, USA), and samples were prepared at a final concentration of 2 μg μl^-1^ in SDS– polyacrylamide gel electrophoresis (PAGE) loading buffer (250 mM Tris-HCl, 20% glycerol, 4% SDS, 0.001% bromophenol blue and 10% β-mercaptoethanol), followed by boiling for 5 min and storage at - 20ºC until use. Ten to 30 μg of each sample were loaded onto 10% SDS-PAGE gels (TGX Stain-Free FastCast system, Bio-Rad), and ran at 230 V. Proteins were transferred to nitrocellulose membranes (Schleicher and Schuell, PerkinElmer, Waltham, MA, USA), incubated in 10% non-fat milk diluted in phosphate-buffered saline (PBS) with 0.1% Tween-20 (PBS-T), washed 3x for 10 min each in PBS-T, and then incubated overnight with primary antibodies overnight at 4°C under constant mild agitation. On the next day, membranes were washed again 3x for 10 min each in PBS-T and incubated for 2 h at room temperature with species-specific horseradish peroxidase-conjugated secondary antibodies. **Table S2** lists all primary and secondary antibodies used in this study. Chemiluminescent reactions were performed with the Pierce ECL Western Blotting Substrate kit (Thermo Fisher Scientific, Waltham, USA) and detected using the ChemiDoc Touch Imaging System (Bio-Rad). Bands were quantified with the Image Lab software v. 6.0 (Bio-Rad) and protein levels were expressed as relative to GAPDH to account for loading differences between conditions.

### Filter trap assay

Detection of polyQ aggregates by filter trap assay (FTA) was performed as described previously ^65^, with a modified lysis protocol that substantially improved assay reproducibility (see below). Three hundred thousand cells per well were seeded in 6-wells plated coated with poly-L-lysine (PLL; 0.001% v/v) and 24 h later transfected with 0.15 μg Q71-GFP and 0.9 μg mRFP or one of the V5-DNAJB6b variants (except for the specific conditions in which 0.45 μg V5-DNAJB6b^WT^ was used, as indicated in the appropriate figures) using 6 μg/ well polyethylenimine (PEI) as the transfection reagent. After 48 h, cells were washed twice in cold PBS and scraped in 200 μl lysis buffer [100 mM NaCl, 50 mM Tris-HCl pH7.4, 1 mM MgCl_2_, 0.5% SDS, benzonase nuclease (0.15 U μl^-1^; Merck, Kenilworth, USA), and 1x EDTA-free cOmplete protease inhibitor cocktail (Merck)]. Samples were vortexed for 20 s and incubated on ice for 30 min (5 s vortex every 10 min). Subsequently, the final SDS concentration was adjusted to 2% and samples were stored at -80ºC until further use. Serial dilutions (1.2, 0.24, and 0.048 μg/μl) of each sample were prepared in FTA buffer (150 mM NaCl, 10 mM Tris-HCl pH8.0, 2% SDS, 50 mM dithiothreitol), boiled for 5 min and stored at -20ºC. One hundred μl of each sample were used for FTA.

### His-ubiquitin denaturing pulldown

For the pulldown of mono-/ poly-ubiquitylated polyQ proteins, 3×10^6^ cells/ dish were seeded on 10 cm dishes coated with PLL. On the following day, cells were transfected with 0.6 μg Q71-GFP, 1.2 μg His-Ub, and 5.4 μg V5-DNAJB6b^WT^ or mRFP constructs using 36 μg/ dish PEI as the transfection reagent. Approximately 36 h after transfection, cells were washed twice in collection buffer [1x PBS, 10 mM N-ethylmaleimide (NEM)], scrapped in 400 μl collection buffer, and spun down at 150 g for 5 min at 4ºC. The supernatant was discarded, cells were snap frozen in liquid nitrogen, and stored at -80ºC until further processing. Cells from two 10 cm dishes per condition were then pooled together, lysed in 1 ml lysis buffer (6 M guanidium-HCl, 100 mM Na_2_HPO_4_-NaH_2_PO_4_ pH7.4, 10 mM Tri-HCl pH8.0, 10 mM NEM, 10 mM β-mercaptoethanol, 5 mM imidazole), and sonicated on ice for 5 s at 50% maximum output. Lysates were centrifuged twice at 20,000 g for 30 min at 4ºC with the supernatant being collected into new tubes at each step. Subsequently, 30 μl of each sample were kept as the input fraction and immediately boiled in 4x SDS-PAGE loading buffer. The remaining lysate was then transferred to 15 ml tubes containing 5 ml lysis buffer and 150 μl Ni-NTA agarose beads (Qiagen, Venlo, Netherlands) and incubated for 4 h at room temperature under constant mild agitation. Beads were sequentially washed once for 5 min each with 750 μl of the following buffers: lysis buffer without imidazole, buffer A (8 M urea, 100 mM, 100 mM Na_2_HPO_4_-NaH_2_PO_4_ pH7.4, 10 mM Tri-HCl pH8.0, 10 mM β-mercaptoethanol), buffer B (same as A, but with Tris-HCl pH6.3 instead of pH8.0) with 0.2% triton X-100 (TX-100), and buffer B with 0.1% TX-100. Mono-/ poly-ubiquitylated species were recovered from beads by a 20 min incubation in 75 μl elution buffer (200 mM imidazole, 150 mM Tris-HCl pH6.7, 30% glycerol, 5% SDS, 720 mM β-mercaptoethanol) followed by boiling for 5 min with SDS-PAGE loading buffer. Input and pulldown samples were then analyzed by Western blot with anti-GFP and anti-ubiquitin antibodies.

### Immunofluorescence

For microscopic analysis, 2×10^5^ cells were seeded on coverslips coated with PLL and cultured for 48 h. Cells were washed twice with PBS, fixed for 15 min in 4% paraformaldehyde, washed twice with PBS for 5 min each, and permeabilized for 5 min with PBS plus 0.1% TX-100. After an additional incubation in PBS for 5 min, primary antibodies (see **Table S2**) diluted in PBS+ [1x PBS, 0.5% bovine serum albumin (w/v), 0.15% glycine (w/v)] were added and cells were incubated overnight at 4ºC. On the following day, cells were washed four times with PBS+ and incubated with fluorescently labelled secondary antibodies (see **Table S2**) diluted in PBS+ for 1.5 h at room temperature. Following two last washes with PBS+, nuclei were stained with Hoechst 33342 (Thermo Fisher Scientific) for 10 min. Coverslips were mounted on microscopy slides with 80% glycerin, sealed, and analyzed on a Leica DM6 B fluorescence microscope equipped with a 63x/1.3 NA oil immersion objective (Leica Microsystems, Buffalo Grove, USA). Images with same intensity were processed and analyzed using the FIJI ImageJ v.2.1.0/1.53c software. For calculation of the nucleus: cytosol ratio of ubiquitylated proteins, fluorescence intensities were estimated in regions of interest of equal area in both the nucleus and cytoplasm of the same cell.

### Live cell imaging

Time course analysis of polyQ aggregation and plate confluence were performed using the Incucyte S3 automated microscope (Essen BioScience, Ann Arbor, USA) coupled with a 20x/0.45 objective. Twenty-four hours after seeding 1×10^5^ cells/ well in 24-wells plates, cells were transfected with 0.5 μg FRT-TO empty vector or V5-DNAJB6b^WT^, or co-transfected with 0.075 μg Q71-GFP plus 0.5 μg mRFP or V5-DNAJB6b^WT^, always in the presence of 3 μg PEI. Brightfield and fluorescence (440-480 nm excitation, 504-544 nm emission) images were acquired every 6 h from three technical replicates per biological replicate. Mean confluence values were determined from a custom mask on the brightfield channel using the Incucyte v.2019B-Rev2 analysis software. For the quantification of polyQ puncta, automatically processed images from the green channel were exported and batch-processed via the Spot Detector plugin from the Icy Bioimage Analysis v.2.1.4.0 and R v.4.0.3 applications using the following parameters (optimized through manual inspection): bright spot over dark background, scale 2 (3 pixels) = 55, scale 3 (7 pixels) = 50, range of accepted objects (in pixels) = 10 to 1000, maximum intensity > 190, surface < 100.

For FLIP, FRAP and condensate frequency analysis, 3×10^5^ cells were seeded in 35 mm glass bottom dishes covered with poly-D-lysine (MatTek Corporation, Ashland, USA) and co-transfected 24 h later with 0.15 μg Q71-GFP plus either 0.9 μg mRFP, mCherry, V5-DNAJB6b^WT^, or mCherry-DNAJB6b^WT^ constructs using Lipofectamine 2000 Reagent (Thermo Fisher Scientific). Cells were analyzed on a Zeiss LSM 780 laser scanning confocal microscope coupled to a XL S1 Dark incubation chamber (Carl Zeiss Microscopy, Jena, Germany), with a 63x/1.3 NA glycerin immersion objective. For FRAP and FLIP-like analyses, rectangular regions of interest (ROIs) of same height and similar area were bleached with a 488 nm laser at maximum power, and recovery/ loss of fluorescence intensity were recorded within ROIs only (FRAP), or across the whole cell (FLIP-like, continuous bleaching to reveal putative condensates). Data were acquired at maximum speed with the Zen Black software (Carl Zeiss Microscopy). For FRAP, 100 pre-bleaching images were recorded, followed by two consecutive bleaching pulses and 900 post-bleaching images (average 0.03 s per image; approximately 27.2 s in total). Fluorescence recovery profiles were normalized to the average pre-bleaching values, excluding the first 50 recordings to account for measurement fluctuations, and data were fitted to a two-phase association model using GraphPad Prism v.9.1.2 to estimate half-lives and immobile fractions. For FLIP-like condensate analysis, 10 pre-bleaching and 90 consecutive bleaching images of whole cells were recorded (average 0.97 s per image; approximately 1.6 min in total).

---

**Table S1:**
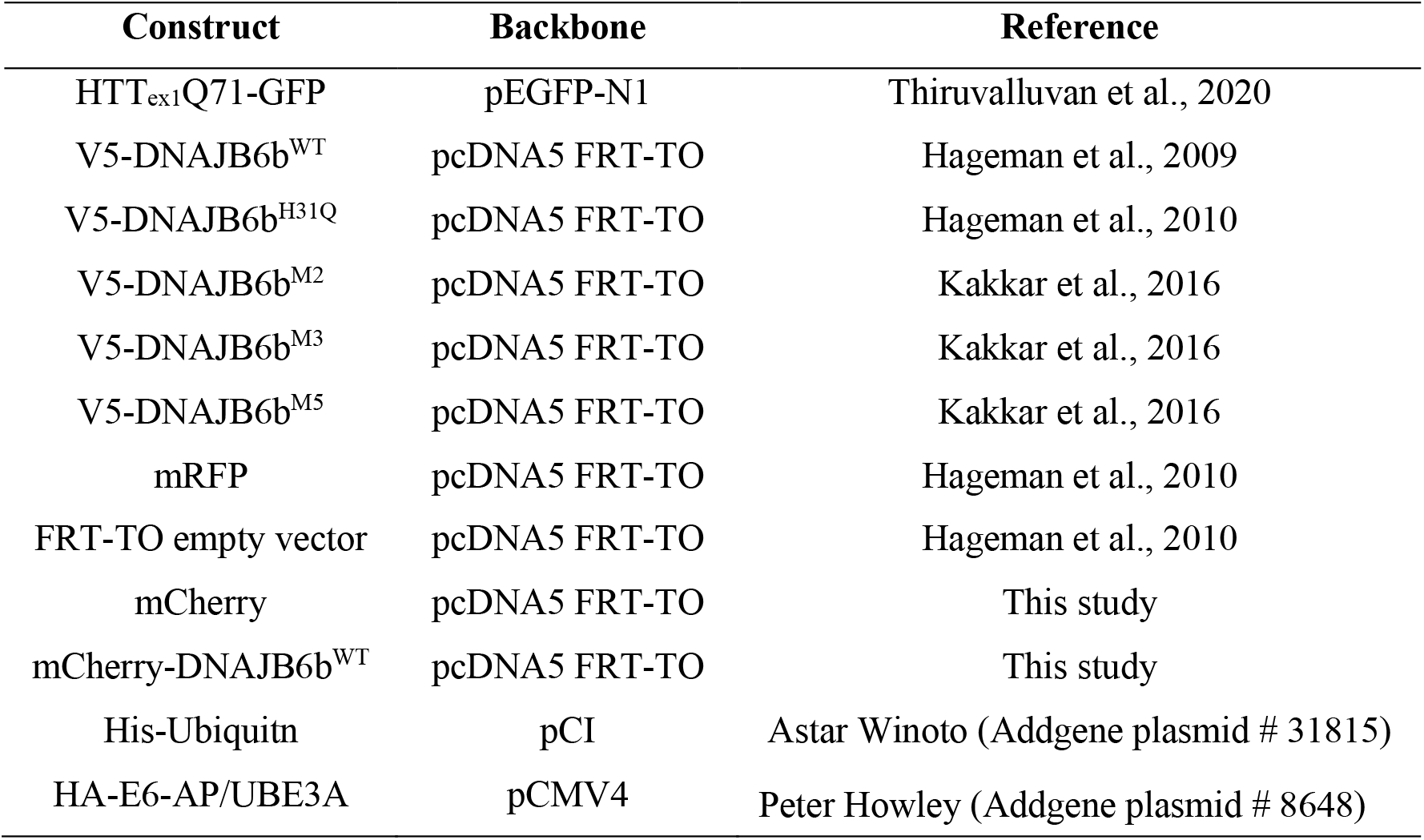
DNA constructs used in the present study.

| **Construct** | **Backbone** | **Reference** |
| --- | --- | --- |
| HTT<sub>ex1</sub>Q71-GFP | pEGFP-N1 | Thiruvalluvan et al., 2020 |
| V5-DNAJB6b<sup>WT</sup> | pcDNA5 FRT-TO | Hageman et al., 2009 |
| V5-DNAJB6b<sup>H31Q</sup> | pcDNA5 FRT-TO | Hageman et al., 2010 |
| V5-DNAJB6b<sup>M2</sup> | pcDNA5 FRT-TO | Kakkar et al., 2016 |
| V5-DNAJB6b<sup>M3</sup> | pcDNA5 FRT-TO | Kakkar et al., 2016 |
| V5-DNAJB6b<sup>M5</sup> | pcDNA5 FRT-TO | Kakkar et al., 2016 |
| mRFP | pcDNA5 FRT-TO | Hageman et al., 2010 |
| FRT-TO empty vector | pcDNA5 FRT-TO | Hageman et al., 2010 |
| mCherry | pcDNA5 FRT-TO | This study |
| mCherry-DNAJB6b<sup>WT</sup> | pcDNA5 FRT-TO | This study |
| His-Ubiquitin | pCI | Astar Winoto (Addgene plasmid # 31815) |
| HA-E6-AP/UBE3A | pCMV4 | Peter Howley (Addgene plasmid # 8648) |

**Table S2.** Primary and secondary antibodies used in the present study.

| **Antibody** | **Company** | **Catalog Number** | **Raised in (clonality)** | **Primary/secondary** | **Dilution (purpose)** |
| --- | --- | --- | --- | --- | --- |
| BAG1 | Cell Signaling | 3920 (3.10G3E2) | Mouse (monoclonal) | Primary | 1:1,000 (WB) |
| BAG3 | Abcam | ab47124 | Rabbit (polyclonal) | Primary | 1:1,000 (WB) |
| DNAJB1 (HSP40) | Enzo | ADI-SPA-400-D | Rabbit (polyclonal) | Primary | 1:1,000 (WB) |
| DNAJB6 | Home-made | NA | Rabbit (polyclonal) | Primary | 1:1,000 (WB) |
| GAPDH | Fitzgerald | 10R-G109a | Mouse (monoclonal) | Primary | 1:10,000 (WB) |
| GFP | Clontech Laboratories | 632381 (JL-8) | Mouse (monoclonal) | Primary | 1:5,000 (WB) |
| HSP90 | StressMarq | SMC-149 (4F3-E8) | Mouse (monoclonal) | Primary | 1:1,000 (WB) |
| HSPA1A | Enzo | ADI-SPA-810-D (C92F3A-5) | Mouse (monoclonal) | Primary | 1:1,000 (WB) |
| HSPA8 | Enzo | ADI-SPA-815-D (1B5) | Rat (monoclonal) | Primary | 1:1,000 (WB) |
| LC3B/ MAP1LC3B | Novus Bio | NB600-1384 | Rabbit (polyclonal) | Primary | 1:3,000 (WB) |
| P62 | Enzo | BML-PW9860-0025 | Rabbit (polyclonal) | Primary | 1:1,000 (WB, IF) |
| Ubiquitin (K<sup>29</sup>-, K<sup>48</sup>-, and K<sup>63</sup>-linked mono- and poly-Ub chains) | Enzo | BML-PW8810-0100 (FK2) | Mouse (monoclonal) | Primary | 1:1,000 (WB, IF) |
| V5 | Invitrogen | R960-25 | Mouse (monoclonal) | Primary | 1:2,000 (WB) |
| HRP-conjugated anti-mouse | GE Healthcare | NXA931 | Sheep | Secondary | 1:5,000 (WB) |
| HRP-conjugated anti-rabbit | GE Healthcare | NA934 | Donkey | Secondary | 1:5,000 (WB) |
| HRP-conjugated anti-rat | GE Healthcare | NA935 | Goat | Secondary | 1:5,000 (WB) |
| Alexa Fluor 488 anti-mouse IgG (H+L) | ThermoFisher | A-11001 | Goat (polyclonal) | Secondary | 1:1,000 (IF) |
| Alexa Fluor 633 anti-rabbit IgG (H+L) | ThermoFisher | A-21070 | Goat (polyclonal) | Secondary | 1:1,000 (IF) |

HRP: horseradish peroxidase; IF: immunofluorescence; NA: not applicable; WB: Western blot.

**Supplementary Figure 1.**
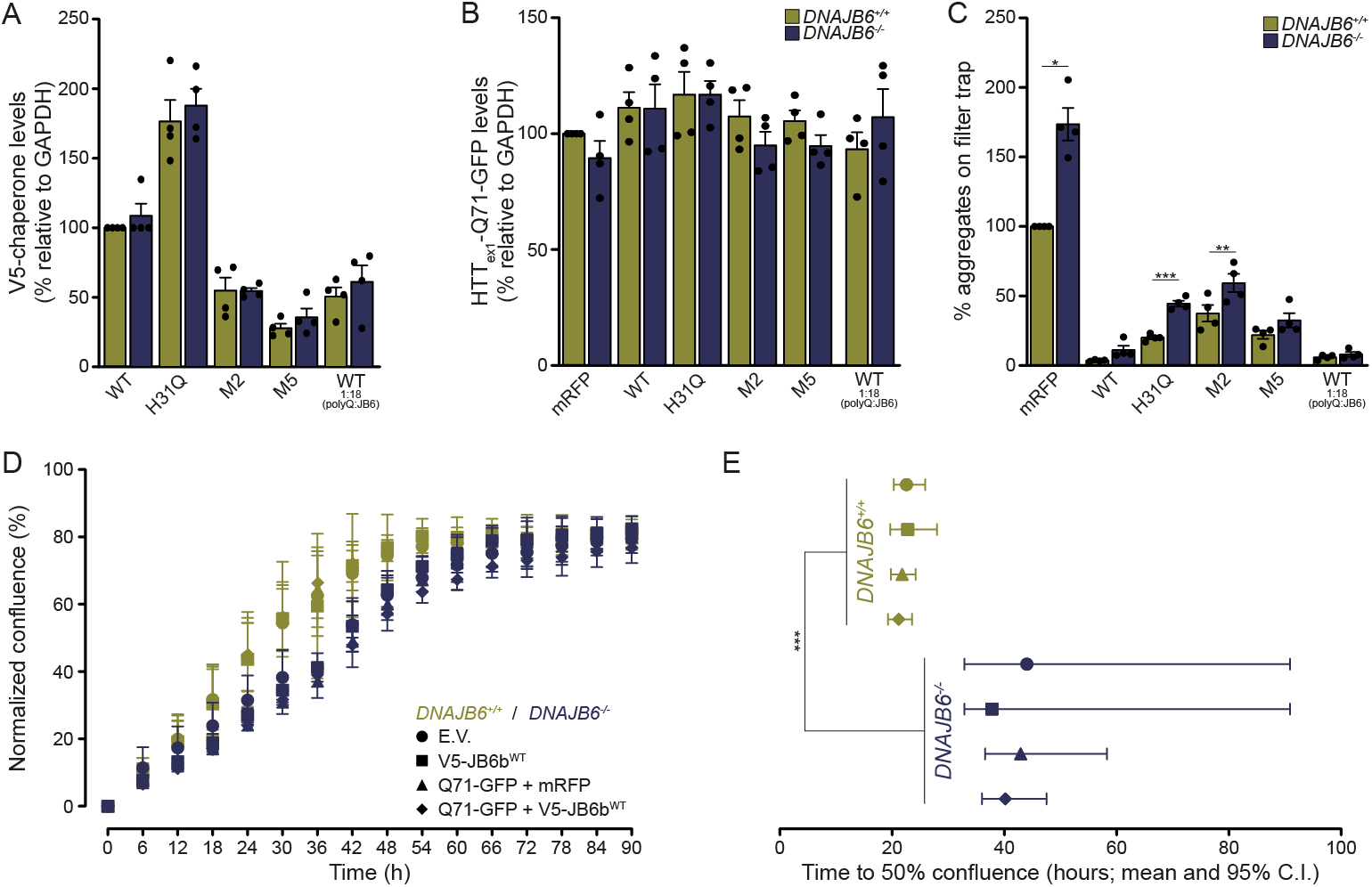
**A (relates to main figure 1D)**: Quantification of distinct V5-tagged DNAJB6b variants used and detected by Western blot. **B (relates to main figure 1D):** Quantification of the ∼70 kDa GFP-positive band detected by Western blot, corresponding to the total steady-state Q71-GFP levels in the running gel. **C (relates to main figure 1F):** Quantification of the percentage of aggregates on filter trap, as shown in main figure 1E, for four independent experimental replicates, and with all variants expressed as relative to the +mRFP only condition in HEK293T wildtype (*DNAJB6*^*+/+*^) cells. * p=0.008, one sample t-test; ** p=0.002, *** p<0.001, one-way ANOVA with Sídák’s multiple comparisons test. **D (relates to main figure 1B):** Quantification of cell confluence via live cell imaging time course in HEK293T wildtype (*DNAJB6*^*+/+*^) or DNAJB6 knockout (*DNAJB6*^*-/-*^) cells co-transfected with Q71-GFP and either mRFP or V5-DNAJB6b. **E (relates to main figure 1B):** Plotting of the time to reach 50% confluence [mean and 95% confidence intervals (C.I.)] for the conditions shown in (D), as estimated by sigmoidal dose-response curves with variable slope fit. *** p<0.001 (F_1,370_=98.74). Error bars represent standard errors of the mean (A-C) or standard deviations (D, E).

**Supplementary Figure 2 (relates to main figures 1G-J).**
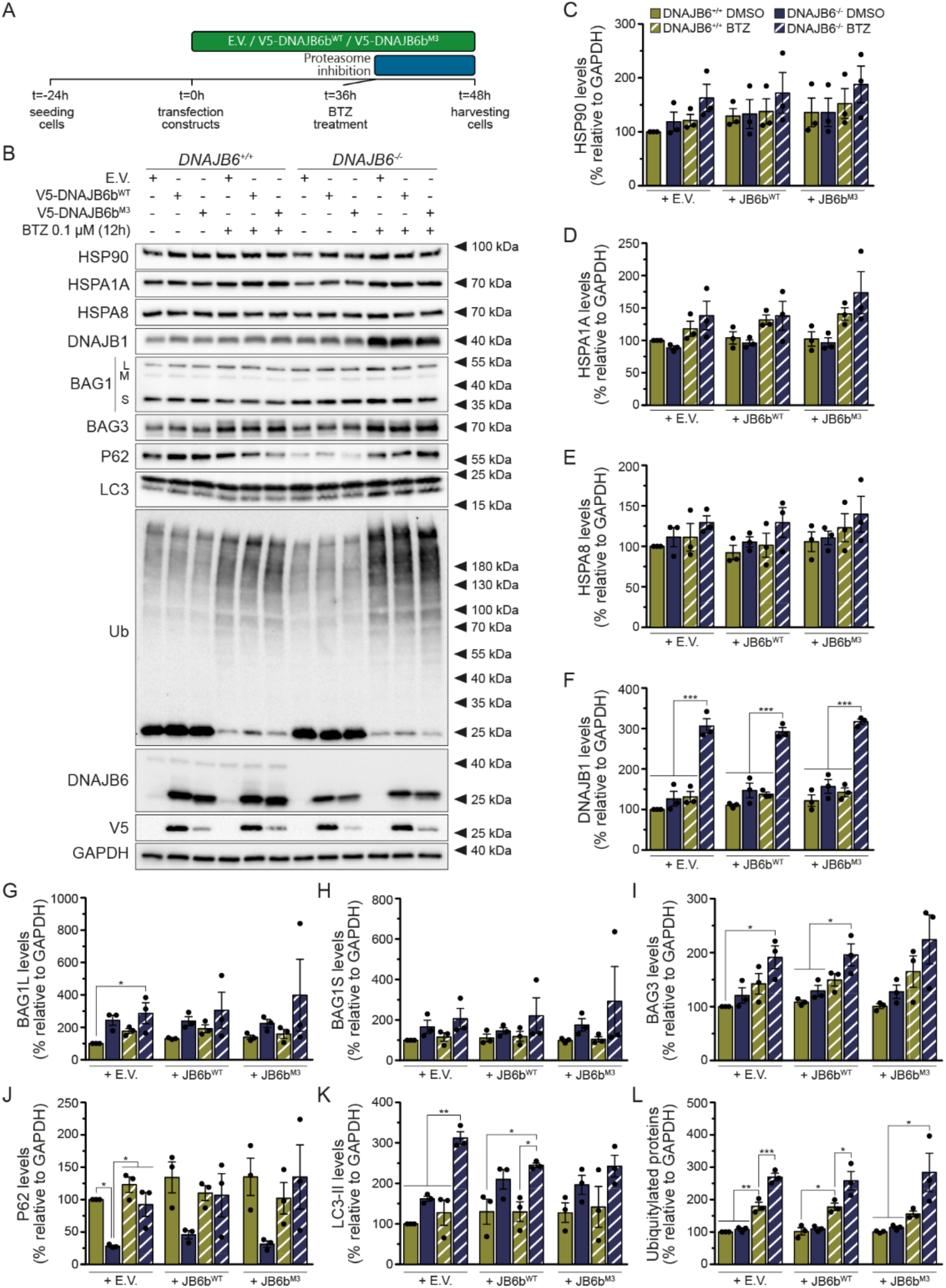
**A:** Diagram illustrating the experimental regimen for the inhibition of proteasomal activity with bortezomib (BTZ) for the last 12 hours before harvesting HEK293T wildtype (*DNAJB6*^*+/+*^) or DNAJB6 knockout (*DNAJB6*^*-/-*^) cells transfected with either empty vector (E.V.), wildtype DNAJB6b (V5-DNAJB6b^WT^), or the polyQ interaction-deficient mutant M3 (V5-DNAJB6b^M3^). **B:** Representative Western blot images of the steady-state levels of selected protein quality control components in cells from both genotypes treated as indicated. The L, M, and S letters next to the BAG1 image refer to the large, medium, and short BAG1 isoforms, respectively. **C-L:** Quantification of Western blot bands corresponding to HSP90 (C), HSPA1A (D), HSPA8 (E), DNAJB1 (F), BAG1L (G), BAG1S (H), BAG3 (I), P62 (J), LC3-II (K), and total ubiquitylated proteins (L). BAG1M was not quantified due to low signal yield. * p<0.05, ** p<0.01, *** p<0.001 (one-way ANOVA with Sídák’s multiple comparisons test for samples transfected with the same construct). Data are represented as means and standard errors of the mean of three independent experimental replicates.

**Supplementary Figure 3 (relates to main figures 1G-J).**
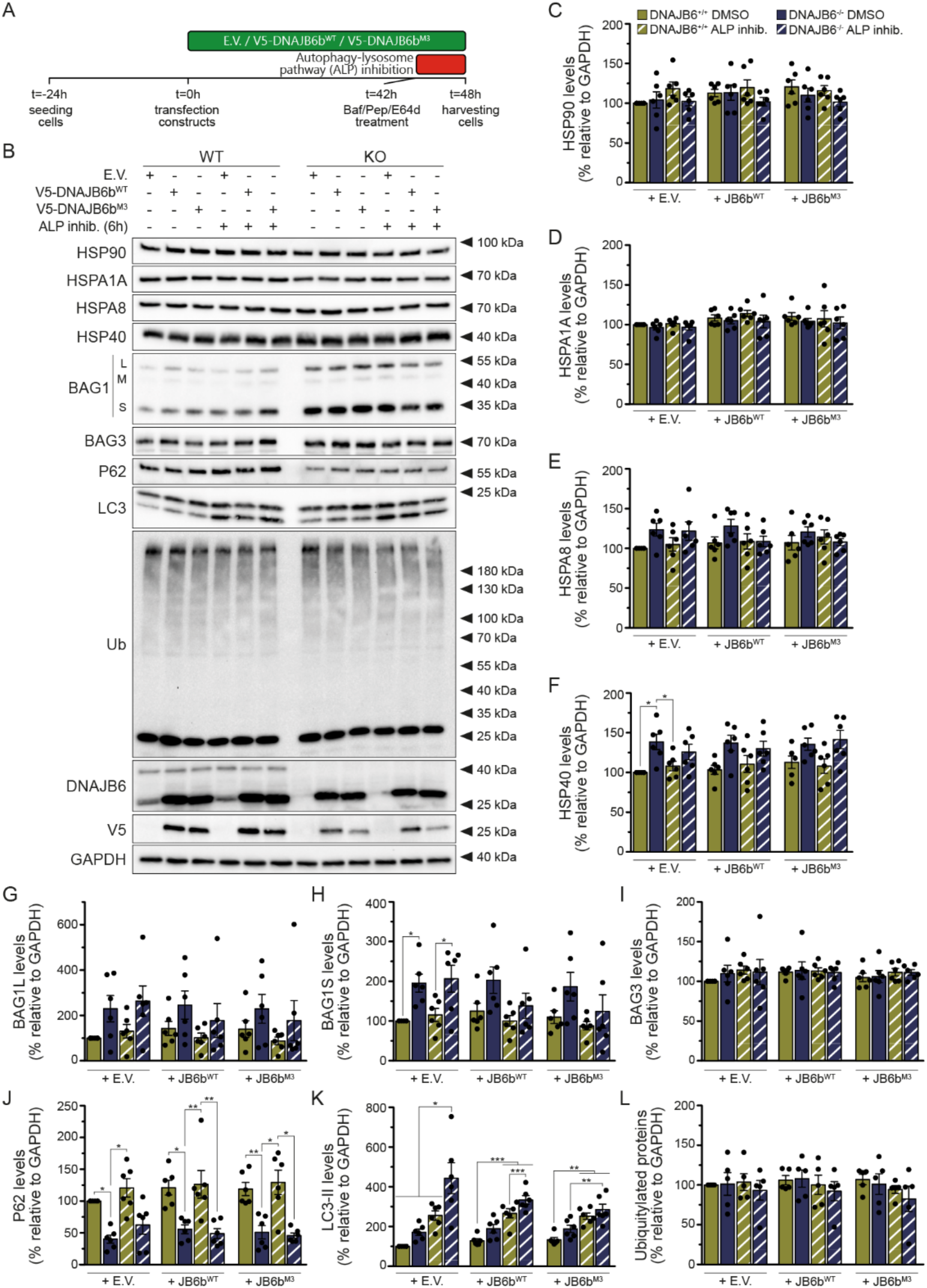
**A:** Diagram illustrating the experimental regimen for the inhibition of autophagic activity with bafilomycin A, pepstatin A, and E64d (Baf/Pep/E64d; ALP inhib.) for the last 6 hours before harvesting HEK293T wildtype (*DNAJB6*^*+/+*^) or DNAJB6 knockout (*DNAJB6*^*-/-*^) cells transfected with either empty vector (E.V.), wildtype DNAJB6b (V5-DNAJB6b^WT^), or the polyQ interaction-deficient mutant M3 (V5-DNAJB6b^M3^). **B:** Representative Western blot images of the steady-state levels of selected protein quality control components in cells from both genotypes treated as indicated. The L, M, and S letters next to the BAG1 image refer to the large, medium and short BAG1 isoforms, respectively. **C-L:** Quantification of Western blot bands corresponding to HSP90 (C), HSPA1A (D), HSPA8 (E), HSP40 (F), BAG1L (G), BAG1S (H), BAG3 (I), P62 (J), LC3-II (K), and total ubiquitylated proteins (L). BAG1M was not quantified due to low signal yield. * p<0.05, ** p<0.01, *** p<0.001 (one-way ANOVA with Sídák’s multiple comparisons test for samples transfected with the same construct). Data are represented as means and standard errors of the mean of six independent experimental replicates. In B, note that a blank gel column separates cells from distinct genotypes in the BAG1, BAG3, P62, LC3, and Ub blots.

**Supplementary Figure 4 (relates to main figure 1).**
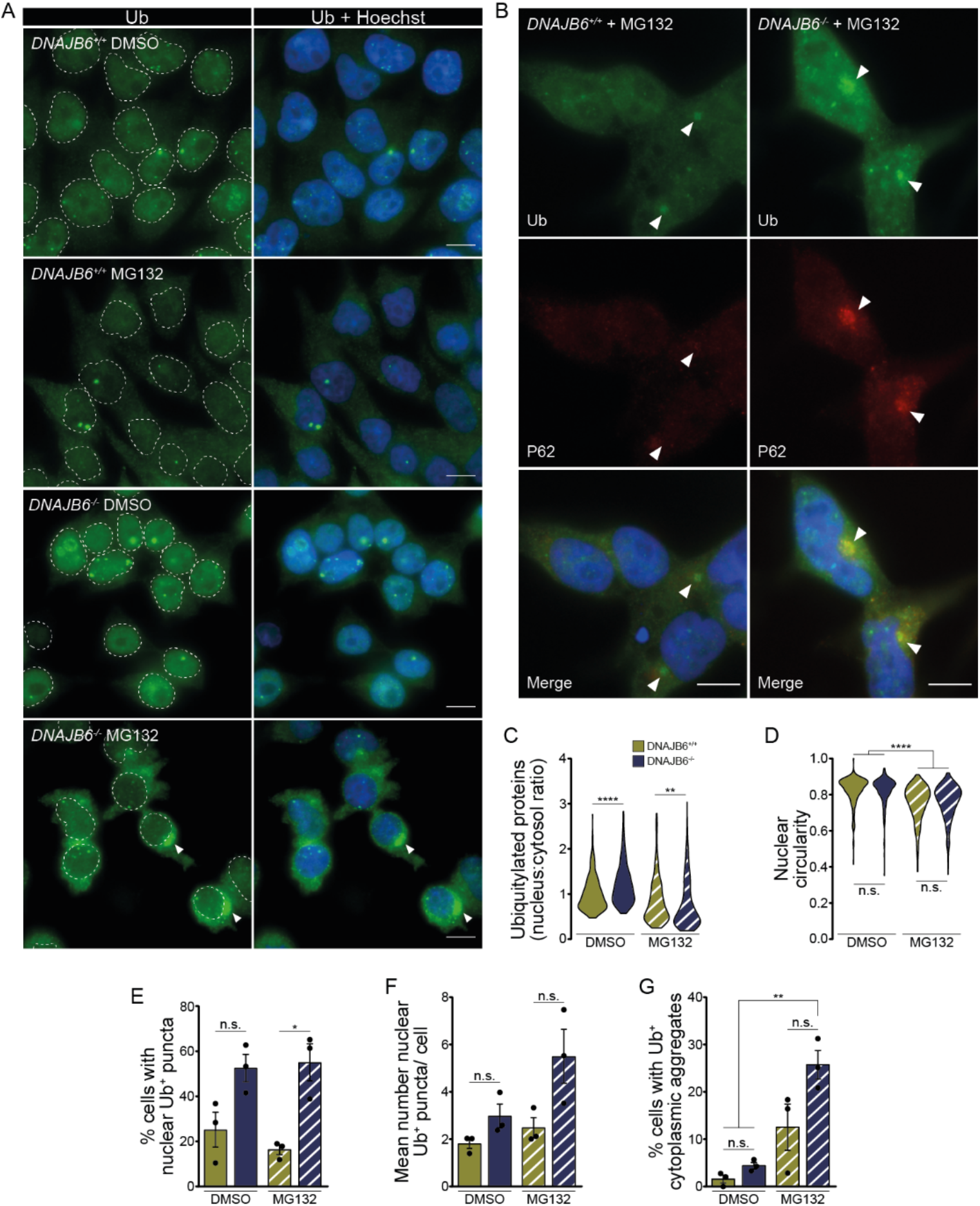
**A:** Representative images of HEK293T wildtype (*DNAJB6*^*+/+*^) or DNAJB6 knockout (*DNAJB6*^*-/-*^) cells treated for 6 hours with either DMSO or the proteasomal inhibitor MG132 and immunostained for ubiquitylated proteins (Ub). Dashed outlines represent the nuclear boundaries, as determined from merged images with the DNA dye Hoechst. Arrowheads: large Ub^+^ deposits. **B:** Representative images of cells from both genotypes treated with MG132 for 6 hours and co-immunostained for Ub (green) and P62 (red). Arrowheads: Ub^+^/P62^+^ foci. Scale bars: 10 μm. **C** and **D:** Violin plots depicting the quantification of the nucleus-to-cytosol ratio of ubiquitylated proteins (C; Kruskal-Wallis with Dunn’s multiple comparisons test; H=318.0, p<0.0001) and nuclear circularity (D; Kruskal-Wallis with Dunn’s multiple comparisons test; H=308.5, p<0.0001) among experimental conditions. Approximately 430 cells per condition across four independent experimental replicates were counted. **E-G:** Quantification of the percentage of cells with nuclear Ub^+^ puncta (E; F_3,8_=8.931, p=0.0062), mean number of nuclear Ub^+^ puncta per cell (F; F_3,8_=5.828, p=0.0207), and percentage of cells with Ub^+^ cytoplasmic aggregate-like structures (G; F_3,8_=13.760, p=0.0016; one-way ANOVA with Tukey’s multiple comparisons test). Data are represented as means and standard errors of the mean from three independent experimental replicates, with approximately 100 cells counted per replicate. n.s.: not significant; * p<0.05, ** p<0.01, **** p<0.0001.

**Supplementary Figure 5.**
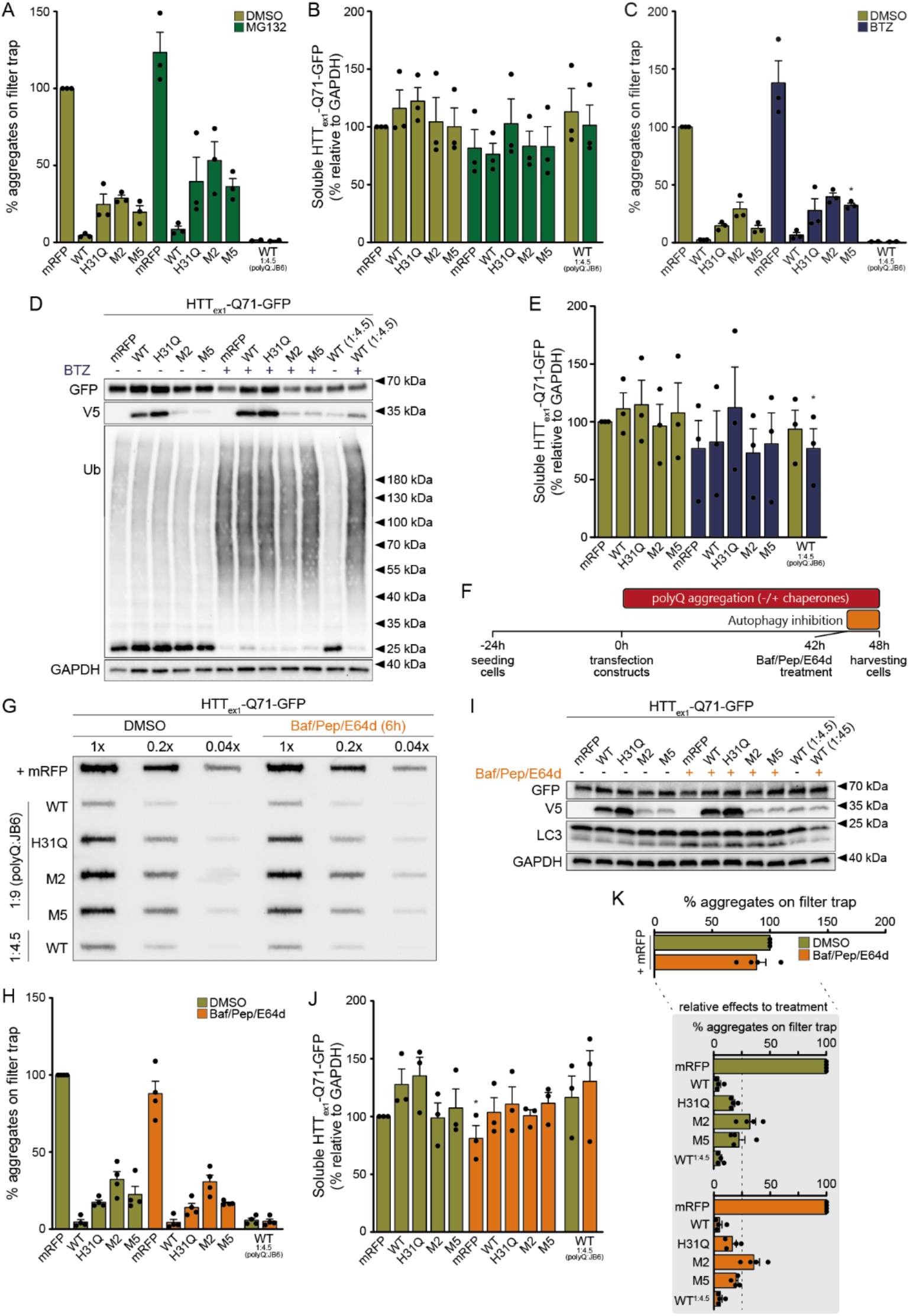
**A and B (relate to main figure 2B-C)**: Quantification of the percentage of aggregates on filter trap (A) and total soluble levels of Q71-GFP (B) in cells co-overexpressing Q71-GFP and mRFP or distinct V5-DNAJB6b variants and treated with either DMSO or MG132 for the last 6 hours before harvesting cells in three independent experimental replicates. **C-E (relate to main figure 2D). C:** Similar to A, but for cells treated with either DMSO or bortezomib (BTZ) for the last 12 hours before harvesting (* p=0.014, ordinary one-way ANOVA with Sídák’s multiple comparisons test). **D:** Representative Western blot images of cells co-overexpressing Q71-GFP and mRFP or distinct V5-DNAJB6b variants and treated with either DMSO or BTZ for the last 12 hours before harvesting. **E:** Quantification of total soluble levels of Q71-GFP in samples as in (D), for three independent experimental replicates. **F:** Diagram illustrating the experimental regimen for the inhibition of autophagy activity with bafilomycin A, pepstatin A, and E64d (Baf/Pep/E64d) for 6 hours before harvesting cells. **G:** Representative filter trap image of cells co-overexpressing Q71-GFP and mRFP or distinct V5-DNAJB6b variants and treated with either DMSO or Baf/Pep/E64d. **H:** Quantification of the percentage of Q71-GFP aggregates on filter trap for samples as in (G) across three independent experimental replicates. **I:** Representative Western blot images of samples as depicted in (G). **J:** Similar to (B) and (E), but for cells treated with either DMSO or Baf/Pep/E64d (* p=0.018, t_df=2_=7.339, one sample t test). **K:** Overview of relative effects of treatment (DMSO vs. Baf/Pep/E64d) in the amount of Q71-GFP aggregates on filter trap across three independent experimental replicates. Quantifications were normalized to the +mRFP control sample within each treatment group. Data are represented as means and standard errors of the mean.

**Supplementary Figure 6 (relates to main figure 2).**
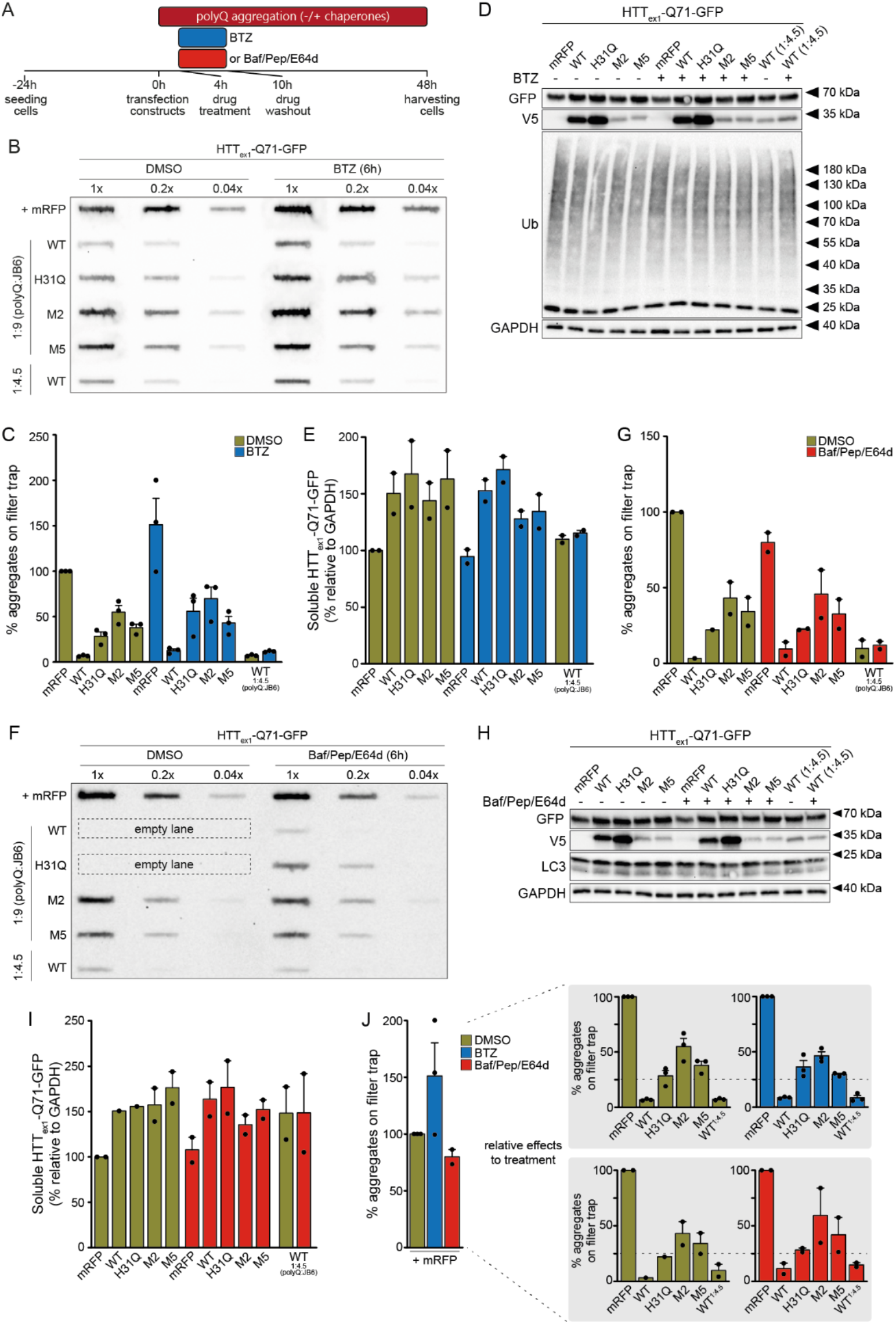
**A:** Diagram illustrating the experimental regimens for the inhibition of proteasomal activity with bortezomib (BTZ) or autophagic activity with bafilomycin A, pepstatin A, and E64d (Baf/Pep/E64d), both for 6 hours immediately after transfection. **B** and **F:** Representative filter trap images of the amount of Q71-GFP aggregates in cells treated with DMSO and either BTZ (B) or Baf/Pep/E64d (F) after co-transfection of Q71-GFP and mRFP or distinct V5-DNAJB6b variants. **C** and **G:** Quantification of the percentage of aggregates on filter trap for the BTZ (C, three experimental replicates) and Baf/Pep/E64d (G, two experimental replicates) regimens. **D** and **H:** Representative Western blot images of samples from the BTZ (D) and Baf/Pep/E64d (H) experimental conditions. **E** and **I:** Quantification of total soluble levels of Q71-GFP in samples from the BTZ (E) and Baf/Pep/E64d (I) regimens (two independent experimental replicates each). **J:** Overview of relative effects of treatment (DMSO vs. BTZ or Baf/Pep/E64d for 6 hours immediately after transfection) in the amount of Q71-GFP aggregates on filter trap across two independent experimental replicates each. Quantifications were normalized to the +mRFP control sample within each treatment group. Data are represented as means and standard errors of the mean.

**Supplementary Figure 7 (relates to main figure 3).**
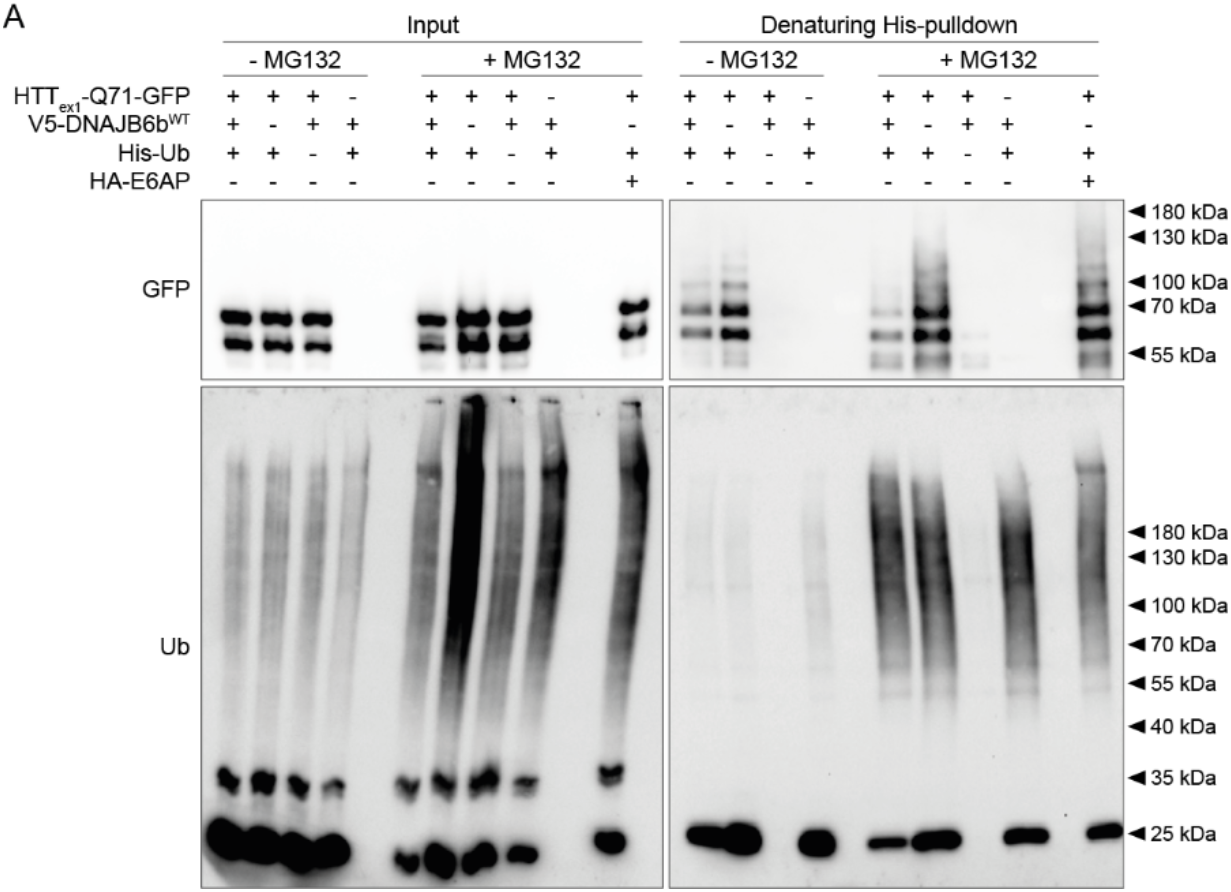
**A:** Original Western blot images (uncropped and not inverted) of data presented on main figure 3. Blots depict denaturing histidine (His)-pulldowns from cells co-transfected with Q71-GFP and either mRFP or V5-DNAJB6b^WT^ in the presence of His-tagged ubiquitin (His-Ub). The ubiquitin E3 ligase E6AP was used as a positive control of ubiquitylation of Q71-GFP.

**Supplementary Figure 8 (relates to main figure 4).**
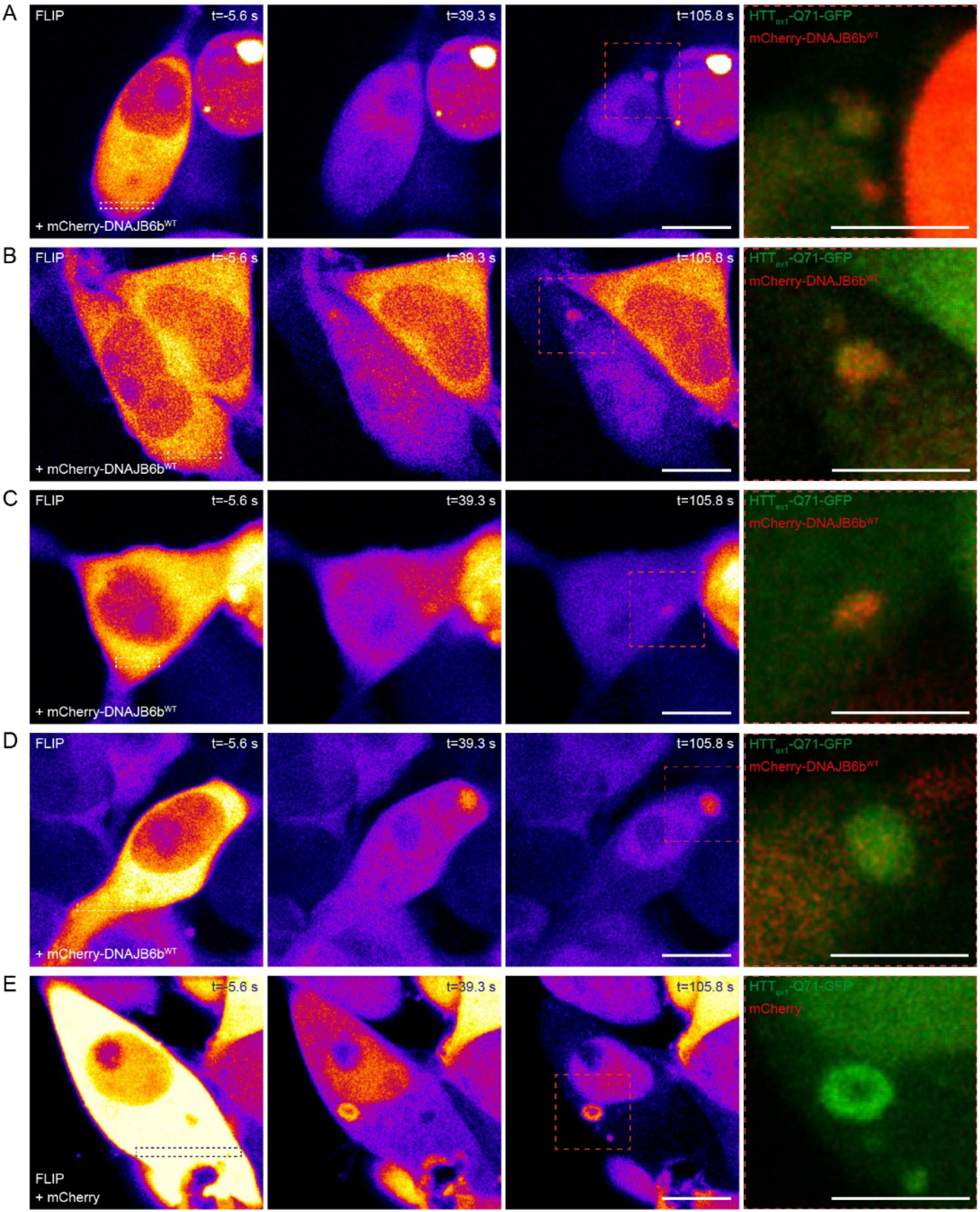
**A-E:** Additional examples of cells co-overexpressing Q71-GFP and either mCherry-DNAJB6b^WT^ (A-D) or mCherry empty vector (E) in which condensate-like polyQ assemblies were identified. False-colored images depict cells immediately before and at selected time points after fluorescence loss in photobleaching (FLIP; dashed rectangular sections). The merged panels to the right represent zoomed sections (dashed orange squares at t=105.8 s) from the GFP and mCherry channels. In all cases, no visible aggregates were observed before FLIP, and internal mobility of the condensate-like structures was confirmed by fluorescence recovery after photobleaching. Scale bars: 10 μm.

